# Modeling the velocity of evolving lineages and predicting dispersal patterns

**DOI:** 10.1101/2024.06.06.597755

**Authors:** Paul Bastide, Pauline Rocu, Johannes Wirtz, Gabriel W. Hassler, François Chevenet, Denis Fargette, Marc A. Suchard, Simon Dellicour, Philippe Lemey, Stéphane Guindon

## Abstract

Accurate estimation of the dispersal velocity or speed of evolving organisms is no mean feat. In fact, existing probabilistic models in phylogeography or spatial population genetics generally do not provide an adequate framework to define velocity in a relevant manner. For instance, the very concept of instantaneous speed simply does not exist under one of the most popular approaches that models the evolution of spatial coordinates as Brownian trajectories running along a phylogeny (Lemey *et al*., 2010). Here, we introduce a new family of models – the so-called “Phylogenetic Integrated Velocity” (PIV) models – that use Gaussian processes to explicitly model the velocity of evolving lineages instead of focusing on the fluctuation of spatial coordinates over time. We describe the properties of these models and show an increased accuracy of velocity estimates compared to previous approaches. Analyses of West Nile virus data in the U.S.A. indicate that PIV models provide sensible predictions of the dispersal of evolving pathogens at a one-year time horizon. These results demonstrate the feasibility and relevance of predictive phylogeography in monitoring epidemics in time and space.

## Introduction

Evaluating the pace at which organisms move in space during the course of evolution is an important endeavor in biology. When considering deep evolutionary time scales, understanding past dispersal events is key to explaining the spatial diversity of contemporaneous species. Over shorter time frames, making sense of the migration patterns of closely related organisms is crucial in building a detailed picture of a population’s demographic past, present and future dynamics. Tracking the spatial dynamics of pathogens during a pandemic, in particular, is of utmost interest as it conveys useful information about the means and the rapidity at which a disease is spreading in a population. Epidemiological data generally consists in records of incidence of the disease at various points in time and space. Yet, estimating the speed at which an organism spreads at the onset of an epidemic from count data is challenging (van den Bosch *et al*., 1992; Tisseuil *et al*., 2016). Similarly, characterizing the migration process from occurrence data in cases where the organism under scrutiny is already well-established in a region is not feasible. These difficulties mainly stem from the fact that count or occurrence data do not convey information about the non-independence between observations due to their shared evolutionary paths.

Genomes carry useful information about the relationships between pathogens. Observed differences between homologous genetic sequences is at the core of phylogenetic and population genetics approaches which provide a sound framework to account for the non-independence between data points in downstream analyses. This framework also accommodates for situations where nucleotide (or protein) sequences are sampled at various points in time (Drummond *et al*., 2003). Heterochronous samples combined with the molecular clock hypothesis (Zuckerkandl and Pauling, 1965) may then serve as a basis to infer the rate at which substitutions accumulate and to reconstruct the time scale of past demographic trajectories of the population under scrutiny (see e.g., (Ho and Shapiro, 2011) for a review).

Designing models for the joint analysis of genetic sequences and their locations of collection was initiated in the middle of the last century by Wright and Malécot who brought forward the isolation by distance model (Wright, 1943; Malécot, 1948). The rise of statistical phylogeography over the last decade proposed alternatives that are less mechanistic but still aim at capturing the main features of the spatial diffusion process. These approaches are also well suited to deal with heterochronous data and handle cases where the population of interest is scattered along a spatial continuum rather than structured into discrete demes. Lemey *et al*. (Lemey *et al*., 2010), in particular, described a hierarchical model whereby spatial coordinates evolve along a phylogenetic tree according to a Brownian diffusion process with branch-specific diffusion rates. The so-called Relaxed Random Walk (RRW) model has since then been used to characterize the spatial dynamics of several pathogens of high public health, societal and agricultural impacts, including Ebola (Dellicour *et al*., 2018b) and the rice yellow mottle (Issaka *et al*., 2021) viruses for instance.

One of the key objectives of the RRW model is to infer the rate at which organisms disperse. Pybus *et al*. (Pybus *et al*., 2012) suggested using a diffusion coefficient which derives from the ratio of the estimated squared displacement between the start and the end of a branch and the corresponding elapsed time. The branch-level ratios are then averaged over the edges in the phylogeny. Pybus *et al*. (Pybus *et al*., 2012) and then Dellicour *et al*. (Dellicour *et al*., 2017) later introduced wavefront-through-time plots, deriving from the displacement between the estimated root location and the most distant tip locations at various points in time. Trovao *et al*. (Trovão *et al*., 2015) considered instead dispersal rates which are defined as ratios of estimated displacements (using great-circle distances) by the elapsed time.

These statistics generally provide a rough characterization of the dispersal process. The limitations of the dispersal statistics mainly stem from the very nature of the RRW model: because Brownian trajectories are nowhere differentiable, the concept of instantaneous speed is simply not defined under that family of models. Also, the sum of displacements deriving from the observation of a Brownian particle at various points in time grows with the square root of the number of (equally spaced in time) observations, making the estimation of an average speed sampling inconsistent. Finally, the analysis of spatial data simulated under the Brownian motion model along birth-death trees shows that the standard dispersal statistics often fail to provide accurate estimates of speed (Dellicour *et al*., 2024).

The present study tackles the issue of dispersal velocity and speed estimation by introducing a new approach that models the instantaneous velocity of lineages explicitly. Under these models, the spatial coordinates of lineages derive from integrating their velocities so that we refer to “Phylogenetic Integrated Velocity” (PIV) models throughout this article. To our knowledge, this study is the first to use the integrated velocity models in a phylogenetic context. Integrated processes are common however in a variety of applications, ranging from population biology (Cumberland and Rohde, 1977) to financial economics (Barndorff-Nielsen and Shephard, 2003). In virology, longitudinal studies measuring CD4 T-cell numbers in cohorts of patients with AIDS have used them to test the hypothesis of “derivative tracking”, in which an individual’s measurements over time tend to maintain the same trajectory (Taylor *et al*., 1994). Closer to phylogeography, integrated processes are instrumental in the field of animal movement ecology (Johnson *et al*., 2008; Hooten and Johnson, 2017). Unlike simple random walks, these processes are not Markovian as the entire track provides information about the next step through the integration. They are thus relevant for accounting for directional persistence. Furthermore, the integrated processes are related to physical models of particles moving on a potential surface (Preisler *et al*., 2013; Russell *et al*., 2018), therefore permitting fine-grained modeling of animal movement telemetry data. One of the goals of the present work is to explore the potential of such approaches in the context of phylogeography, starting with the two simplest and most common models, namely the integrated Brownian and Ornstein-Uhlenbeck processes.

Although velocity is not directly observable from heterochronous and geo-referenced genetic sequences, our results indicate that this quantity can be estimated reliably. Using simulations under realistic spatial population genetics models, we show that the velocity inferred with the new models are more accurate than those deriving from the RRW approach. Velocities estimated from the analysis of multiple West Nile virus data sets were also used to predict the spatial distribution of the pathogen over a one-year time horizon in the U.S.A. Comparison of these predictions to incidence data at the county-level suggests that important features of the spatial dynamics are indeed amenable to reasonably accurate predictions.

Our ability to efficiently monitor and anticipate the spread of emerging epidemics depends on the accuracy with which the pace of dispersal can be quantified. The family of new models introduced in this study provides a relevant tool to achieve this objective. While important aspects of viral evolution may escape prediction indefinitely (Holmes, 2013) and predicting the time and/or location of the next virus outbreak remains out of reach (Wille *et al*., 2021; Holmes *et al*., 2018), the present study shows how predictive phylogeography may complement classical approaches in epidemiology.

## Results

### PIV models: rationale

The main attributes of models that belong to the PIV family are presented first. We focus on the process of interest along a given time interval [0, *t*], corresponding to the length (in calendar time units) of a given branch in the phylogeny of a sample of the organism of interest. Let *X*(*s*) be the random variable representing the location (i.e., the coordinates) of a lineage at time 0 ≤ *s* ≤ *t. Y* (*s*) is its velocity, i.e., the vector that is made of the instantaneous rate at which a lineage changes its position along each dimension of the habitat at time *s*. In all the following, we reserve the term *velocity* for the vector, and *speed* for its scalar norm. Both *X* and *Y* are typically vectors of length two, corresponding to latitude and longitude. The location *X*(*t*) at the end of the branch may then be expressed as follows:

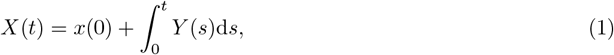

where *x*(0), the location at the time of origin, is fixed. The Brownian Motion (BM) and the RRW models focus on *{X*(*s*), 0 ≤ *s* ≤ *t}*, i.e., the process describing the evolution of the location during a time interval. While, in one dimension, BM models have a single dispersal parameter that applies to all edges in the phylogeny, the RRW model has branch-specific dispersal parameters, in a manner similar to the relaxed clock model (Drummond *et al*., 2006) used in molecular dating.

Instead of modeling the fluctuation of coordinates, PIV models deal with *{Y* (*s*), 0 ≤ *s* ≤ *t}*, i.e., the process describing the variation of velocity in that interval. The dynamics of spatial coordinates then derive from the integration over the velocity as stated in (1) above, hence the name “phylogenetic integrated velocity”. In the following, we introduce two stochastic processes for *{Y* (*s*), 0 ≤ *s* ≤ *t}* and characterize the corresponding distributions of *X*(*t*). In order to simplify the presentation, we provide formulas for univariate processes only in the main text. Formulas for bi-variate (and, more generally, multivariate) processes are given in SI (sections C and D).

### Behavior of PIV models

#### Velocities

The Integrated Brownian Motion (IBM) model relies on a Wiener process with shift and scale parameters *y*(0) and *σ* respectively to model *{Y* (*t*); *t >* 0*}*. That process is Gaussian and we have (Gardiner, 2009):

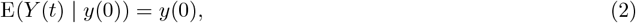

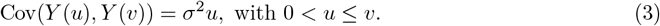

The Integrated Ornstein-Uhlenbeck (IOU) model uses instead a Ornstein-Uhlenbeck (OU) process to describe the evolution of velocity. The mean and variance of velocity at time *t* are given below (Gardiner, 2009):

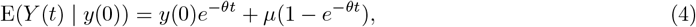

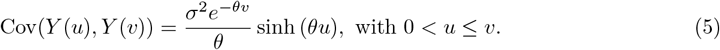

The parameter *θ* in the OU model governs the strength with which *Y* (*t*) is pulled towards the trend *μ*.

#### Spatial coordinates

We now examine the evolution of spatial coordinates under the PIV models. Characterizing the process governing the evolution of spatial coordinates will shed light on the biological relevance of the proposed approach and exhibit the main difference in behavior in comparison with the BM and, by extension, the RRW models. The stochastic processes modeling the fluctuation of velocity being Gaussian, the coordinates also follow a Gaussian process (Cumberland and Rohde, 1977). We give below the mean and variance of the distribution of *X*(*t*) given *x*(0) and *y*(0), the coordinates and velocity at time 0.

When velocity follows a Brownian process (IBM process), we have:

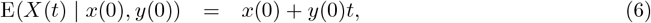

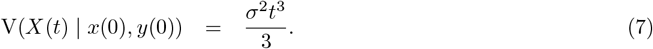

A linear increase of the spatial coordinates is thus expected with a direction that is determined by the initial velocity ((6)). Because of the inertia deriving from their velocity, spatial coordinates of lineages evolving under IBM thus tend to resist changes in their direction of motion, i.e., they exhibit directional persistence (Johnson *et al*., 2008). This mean drift is similar to the directional random walk, used e.g. in (Gill *et al*., 2016) to model the spatial spread of HIV-1. The BM model has a distinct behavior as it authorizes sudden changes of direction. The RRW can even lead to large discontinuous “jumps” from one place to another (Bastide and Didier, 2023). In contrast, the IBM is smoother (differentiable) by design, and well suited to model auto-correlated movements. Moreover, as suggested by (7) above, the variance of coordinates grows cubically in time, thereby allowing the IBM model to accommodate for dispersal events over long distances in short periods of time. This process is thus able to handle fast spatial range expansion, yet with continuous and differentiable trajectories.

The corresponding expectation and variance for the IOU model are given in SI (section A). Here again, the average coordinates at the end of the branch of focus are determined by the coordinates at the start of that branch (*x*(0)) plus the expected displacement (*y*(0)*t*) along that same edge. In this simple IOU model, the velocity of the process converges to the central value *μ*, leading to trajectories with a clear directional trend that are well suited for dispersal along an established spatial gradient. While for small values of *θ* the IOU has a behavior that is similar to the IBM, for larger values of that parameter, its variance grows linearly in time and the process behaves like a directional BM (Gill *et al*., 2016). The auto-correlation (or strength) parameter *θ* is thus interpreted as the amount of directional persistence present in the data (Johnson *et al*., 2008), with small values indicating more dependence to the trajectory path for future moves.

Figure 1 illustrates the behavior of the classical random walk and integrated models along a 5-tip tree. Trajectories of coordinates generated with the BM and Ornstein-Uhlenbeck (OU) versions of the random walk model are intricate, showing abrupt changes of directions in the movements (Fig. 1b, c). The same behavior is displayed by the velocity trajectories under the IBM and IOU models (Fig. 1d, f) as the models are here identical to that used for the BM and OU models indeed. Yet, integrating over these rugged paths gives smooth (differentiable) trajectories of coordinates under the corresponding models (Fig. 1e, g), with particles moving swiftly away from their initial points, illustrating the cubic variance pointed above. The IOU model presented here converges to a (1,1) velocity so that the coordinates of the five lineages show a clear directionality, stronger than that obtained with the OU model (Fig. 1g vs. c).

**Figure 1:**
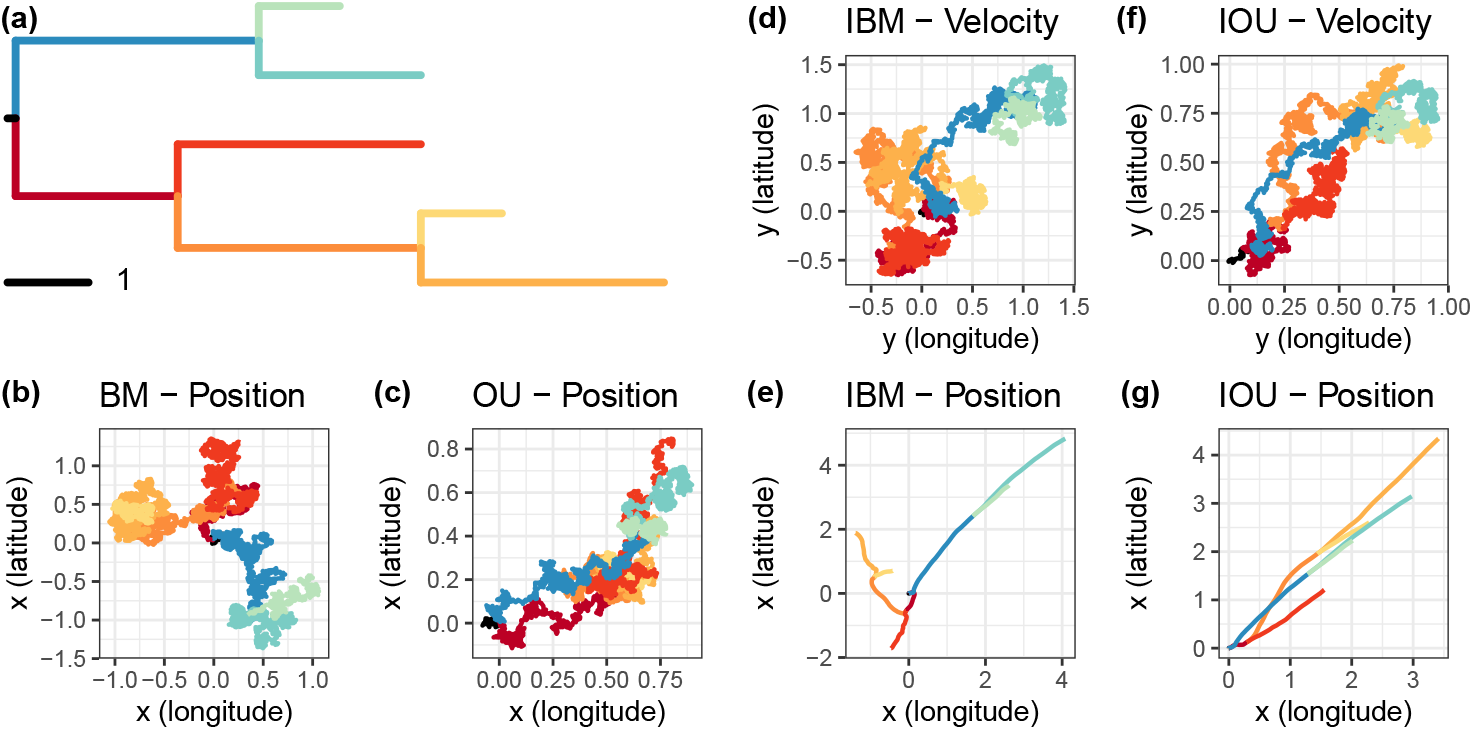
Simulated trajectories of classical random walk and PIV models on a simple tree. Each process was simulated on the tree **(a)**, with branches of matching colors. Movements along the latitude and longitude axes were simulated independently. **(b)** Random walk using Brownian motion (BM) with variance *σ*^2^ = 0.1, and starting point *x*(0) = (0, 0). **(c)** Ornstein-Uhlenbeck process (OU) with stationary variance of *σ*^2^*/*(2*θ*) = 0.1, strength *θ* = 0.17, starting at *x*(0) = (0, 0), and converging to its central value *μ* = (1, 1). **(d)** and **(e)** Velocity *y* and position *x* of an Integrated Brownian Motion (IBM) with variance *σ*^2^ = 0.1, starting point *x*(0) = (0, 0), and starting velocity *y*(0) = (0, 0). **(f)** and **(g)** Velocity *y* and position *x* of an Integrated Ornstein-Uhlenbeck (IOU) with stationary variance *σ*^2^*/*(2*θ*) = 0.1, strength *θ* = 0.17, central trend of *μ* = (1, 1), starting point *x*(0) = (0, 0), and starting velocity *y*(0) = (0, 0).

### Accuracy of speed estimation

Data sets were simulated under the spatial Lambda-Fleming-Viot (SLFV) model (Etheridge, 2008; Barton *et al*., 2010) and an agent-based spatially explicit transmission chain simulator which aimed at mimicking outbreaks of the Ebola virus in West Africa (Lequime *et al*., 2020). 100 data sets were analyzed for each of these two simulation settings. As traditional speed statistics are typically computed over the whole tree (Dellicour *et al*., 2020a), we assessed the ability of PIV models to estimate tree-level speed by averaging node-level velocities across the tree. The classical “weighted lineage dispersal velocity”(WLDV) (Dellicour *et al*., 2016) was used instead for all analyses performed under the RRW model. As shown recently (Dellicour *et al*., 2024), we expect the WLDV statistic on RRW models to perform poorly, and would like to asses the ability of PIV models to provide more accurate speed estimates.

Examination of the estimated vs. true speed relationship (Fig. 2) indicates that the RRW model systematically underestimates speed and the bias worsens with increasing speed. This bias is strong with data simulated under the SLFV (Fig. 2a) and milder with the Ebola data sets (Fig. 2b), which is expected since the transmission trees generated in the latter case are sampled in time and not ultrametric, making the temporal signal to estimate speed stronger. Nonetheless, true speed values are, on average, 1.6 times larger than those estimated with the RRW for the Ebola data and 22 times larger for the SLFV data (the ratios for IBM are 1.2 for both simulation settings). The SLFV model assumes a finite-size habitat (a square here) and boundary effects, which occur for large and small dispersal values, are expected to impact the estimation of speed under models that ignore this constraint. Yet, the IBM model is largely immune to this issue. While the IOU model underestimates speed for the SLFV data sets, its estimates are less biased than those deriving from the RRW model. The IOU model also tends to overestimate speed on the Ebola data sets. Further examination of these results shows a clear influence of the prior distribution on the strength parameter in the IOU model, a phenomenon already observed in (Cornuault, 2022).

**Figure 2:**
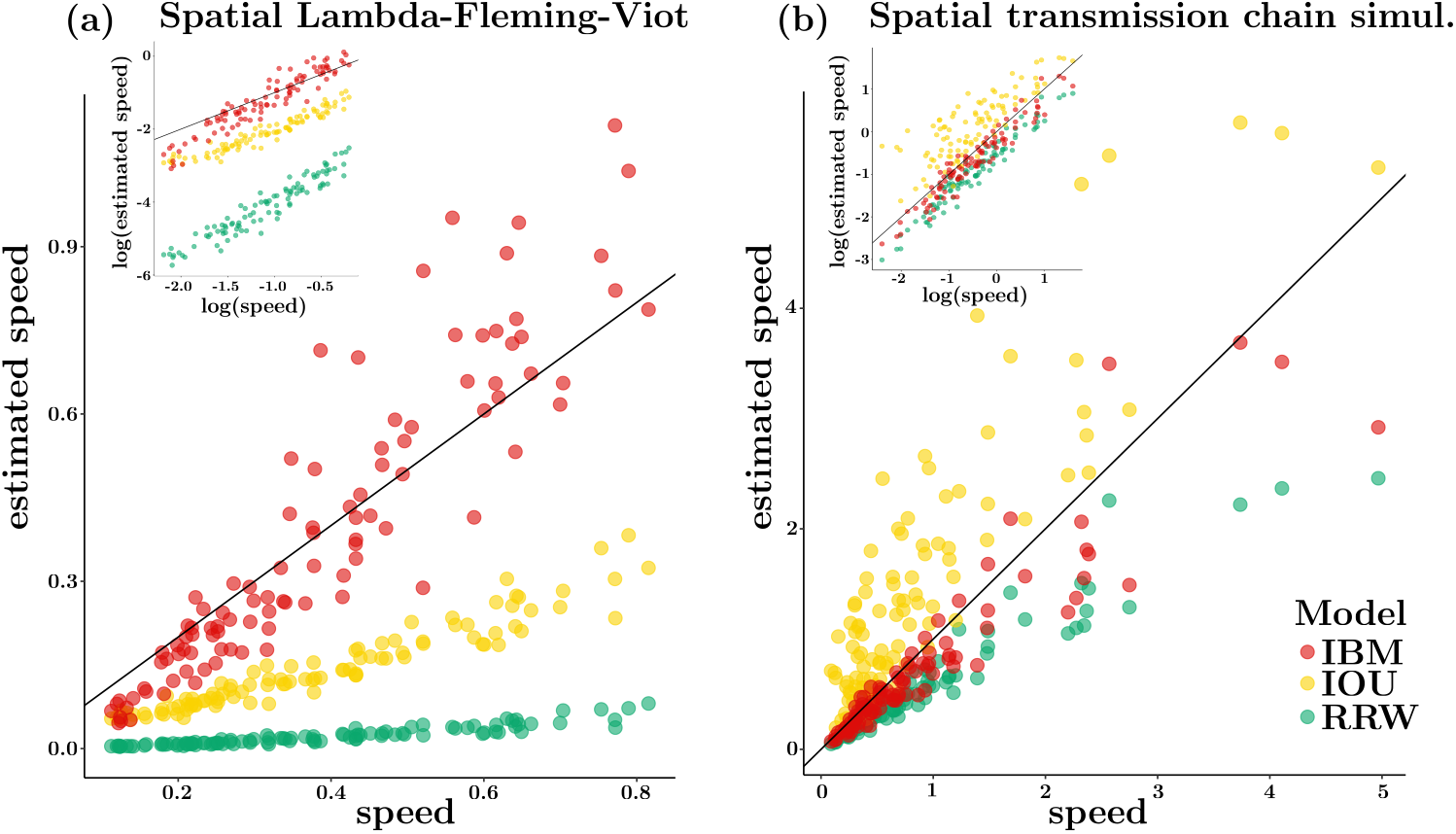
Accuracy of speed estimation under the RRW and PIV models. True (*x*-axis) vs. estimated (*y*-axis) speed. Estimates were obtained under the IBM, IOU and RRW models. (a) 100 data sets were simulated using the spatial Lambda-Fleming-Viot process on a 10 by 10 square. (b) 100 data sets were simulated under a random walk model inspired by the Ebola epidemic in West Africa (see main text). The insets give the log-log scatterplots of the estimated vs. true speeds. The *y* = *x* line is shown in black on each plot.

### Dispersal dynamics of the West Nile virus in the U.S.A

The phylogeography of the WNV in the U.S.A. has been studied extensively (see, e.g., (Dellicour *et al*., 2020a)). The origin of this epidemic took place in New York City during the summer 1999 (Lanciotti *et al*., 1999; Campbell *et al*., 2002). By 2004, human infections, veterinary disease cases or infections in mosquitoes, birds, or sentinel animals had been reported to the Centers for Disease Control and Prevention (CDC) in most counties.

We fitted the PIV and RRW models to several subsets of the 801 geo-referenced sequence data set analyzed in (Dellicour *et al*., 2020a). PIV models are less flexible than the RRW approach as they do not authorize sudden changes of direction, as noted earlier (and see SI, section B). Therefore, ensuring that both approaches nonetheless provide comparable fit to the data is a prerequisite to further analyses. We then used the IBM model to predict the dispersal patterns and evaluate these predictions through the comparison with incidence data for the 2000-2007 time period.

#### Model comparison

We compared the fit of the RRW and PIV models to the WNV data using cross-validation of location information. Cross-validation is a powerful model comparison technique in the context of phylogenetic factor analysis (Hassler *et al*., 2022). Using a subset of 150 data points chosen uniformly at random among the 801 available observations, a leave-one-out procedure was applied to the sample coordinates. Each tip location was first hidden and its posterior density was estimated using MCMC from the remaining 149 locations and all 150 sequences (see SI, section G).

Figure 3 shows the distributions of the great circle distances between the observed and reconstructed tip locations as inferred under the PIV and the RRW models, along with that of uniform at random predictions. The three phylogeographic models have similar behavior overall with a majority of distances between true and reconstructed tip locations ranging between 238 km (25% quantile of distribution from MCMC output pooled across models) and 950 km (75% quantile) with a median of 450 km. In contrast, if inferred locations are uniform at random within the U.S.A. (excluding Alaska and Hawaii), the median distance is 1,564 km, i.e., more than three times that estimated with the phylogenetic models. This result demonstrates the ability of these models to extract meaningful signal from the data, even though these approaches do not account for habitat borders (while the uniform predictor does so). Examination of the posterior distribution deriving from each model taken separately indicates that the median distances obtained under the IBM, IOU and RRW models are 474, 496 and 416 km respectively. While the fit of the RRW model is superior to that of the PIV models, the performance of the three models are nonetheless qualitatively similar.

**Figure 3:**
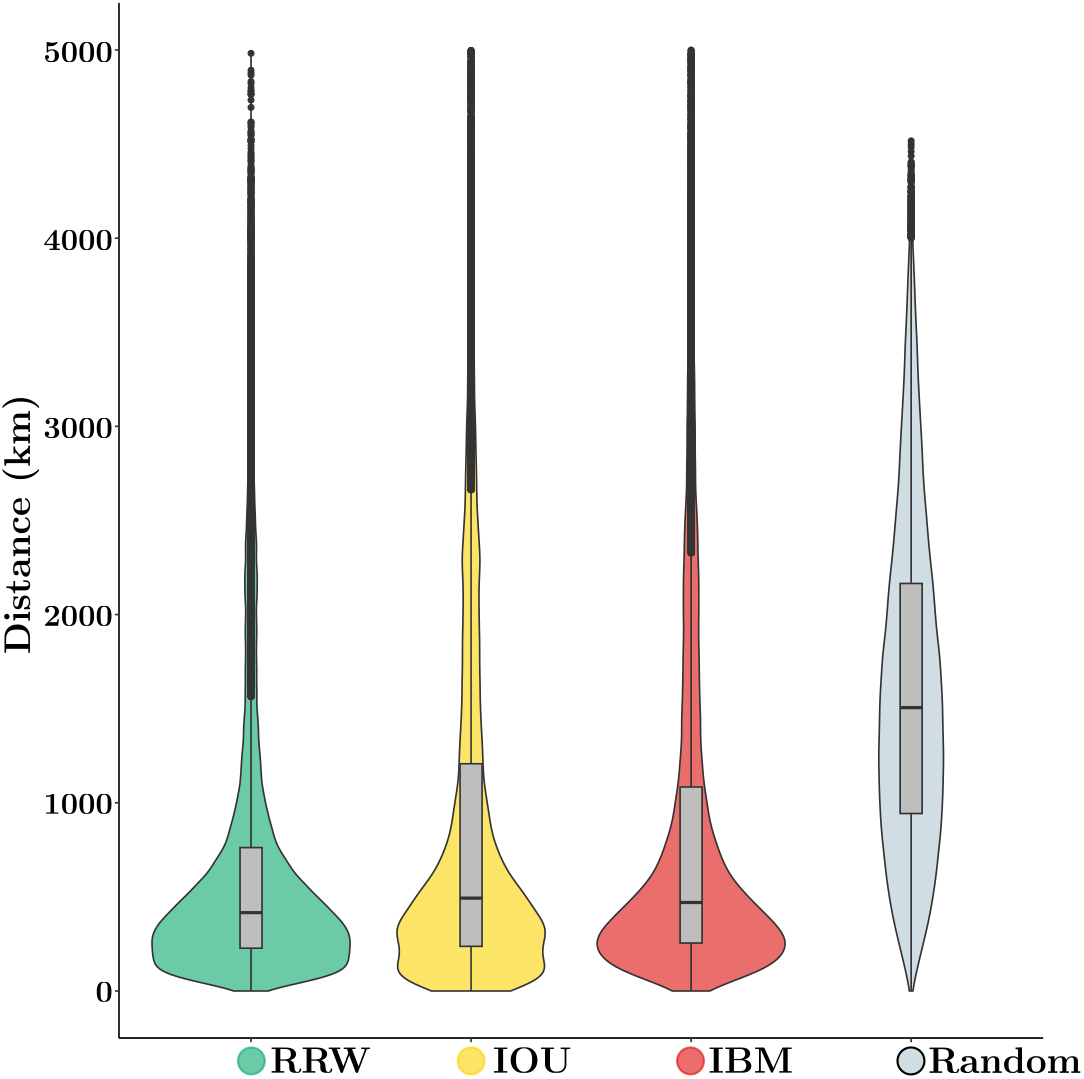
Distribution of the distance between true and estimated tip coordinates under the PIV, the RRW models and uniform at random predictions, WNV data. Cross validation was used to predict the locations of held-one-out tip lineages under the RRW and PIV models. ‘Random’ gives the distance between two locations selected uniformly at random within the U.S.A. The *y* axis gives the great circle distance between coordinates (in km).

#### Predicting dispersal using PIV models

PIV models enable the estimation of dispersal velocity of each sampled lineage. These velocities may then serve as a basis to predict the spatial distribution of the underlying population in the near future. Here, we tested the ability of the IBM model to anticipate the dynamics of dispersal of the WNV in the early and later stages of the epidemic.

Sequences collected earlier than December of year *Y* were randomly subsampled from the complete data set with exponentially increasing weights given to recent samples. Data sets with 150 sequences were obtained except for years 2000-2002 where smaller sample sizes were considered due to a lack of observations in this time period. Estimated posterior distributions of velocities at the tips of the obtained phylogeny under the IBM model were then used as predictors of the spatial distribution of the virus in year *Y* + 1 (see section “Predictive phylogeography” in “Material and Methods”). The predicted occurrences were compared to yearly incidence data collected at the county level.

Figure 4 shows the incidence and the predicted occurrence of the WNV in the early stages in the epidemic. Samples for years 2000, 2001 and 2002 included only 7, 19 and 68 geo-referenced sequences, thereby making any prediction inherently challenging. For instance, predictions for year 2000 are overly dispersed and sensitive to priors (see SI, section H). Also, while the virus had reached Florida by 2001, our model failed to predict its presence south of North Carolina. Predictions for subsequent years rely on larger numbers of observations and demonstrate the relevance of our approach. Indeed, the PIV model successfully predicted the arrival of the pathogen along the west coast of the U.S.A. by the end of 2002. It also correctly predicted that the north west corner of the country would remain largely virus-free until the end of 2003. Predictions deriving from the RRW show qualitatively distinct patterns with a widespread presence of the virus for years 2003 and 2004 that contrasts with incidence data (see SI, section H). Overall, the RRW shows a higher sensitivity (average of 0.89 over all years for the RRW, vs. 0.72 for the IBM), but a lower specificity (average of 0.36 for the RRW, vs. 0.56 for the IBM), consistent with wider and rather vague predicted regions.

**Figure 4:**
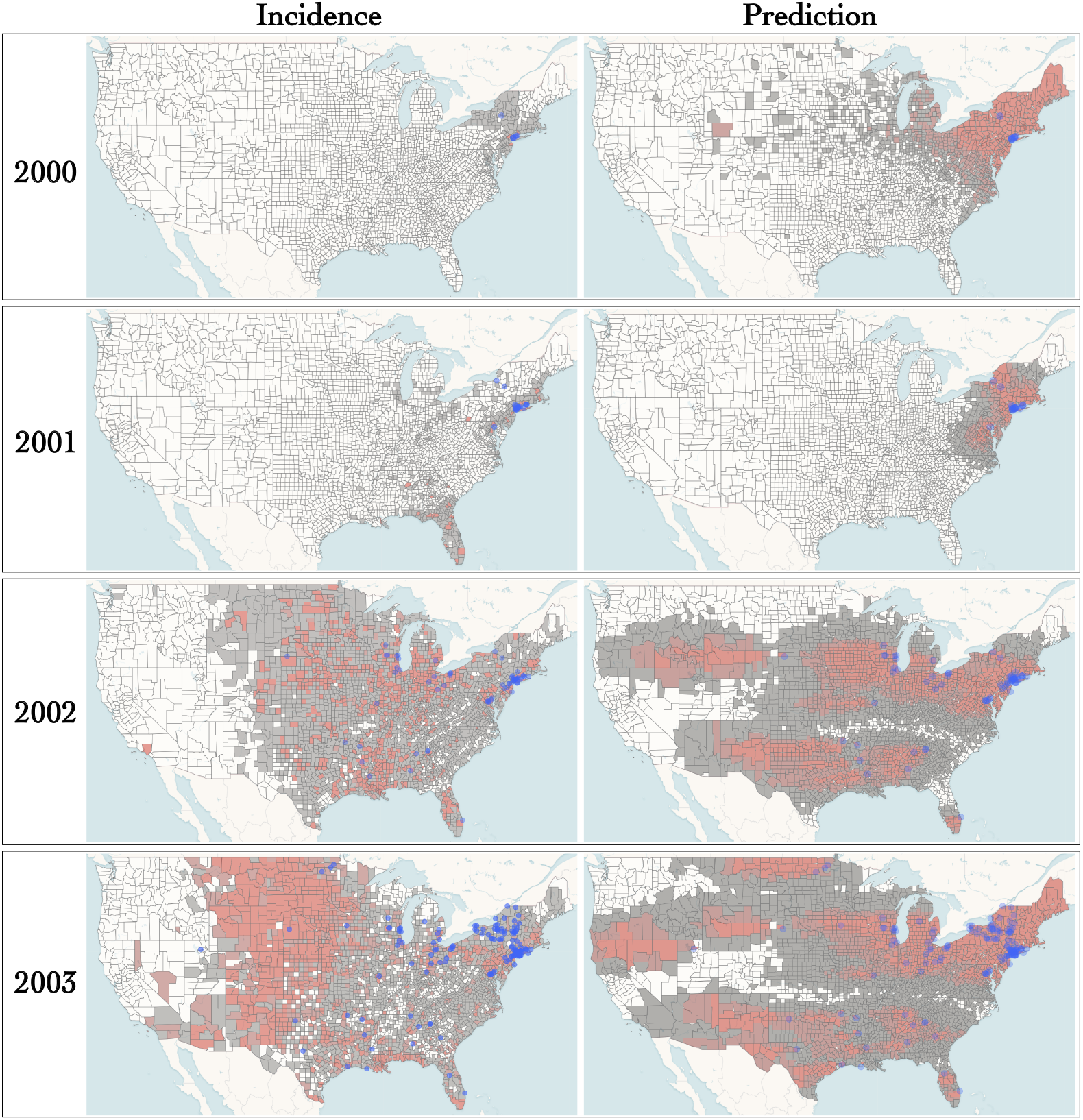
Incidence and predicted occurrence of WNV in the early phase of the epidemic (model for prediction: IBM). Purple dots correspond to sampled locations. Incidence data (left) for each year and each county was obtained from the CDC. For year *Y*, predicted occurrence of the WNV (right) was inferred using data collected earlier than the end of December of year *Y* − 1. The maps were generated with EvoLaps2 (Chevenet *et al*., 2024)

By 2004 the virus reached an endemic state and the spatial dynamics of the epidemic diverged from that of the early stages. Figure 5 shows the results for the 2004-2007 time period. Prediction at local spatial scales has limited accuracy. For instance, a high probability of occurrence was systematically estimated for the states in the North East corner of the country and the south of Texas while incidence was generally mild in these areas. Note that the difference between predicted and observed occurrence could reflect a relatively lower ecological suitability of these regions to host local WNV circulation, thereby serving a useful purpose. Moreover, the IBM model correctly predicts the expansion of the epidemic north of California and Nevada between 2005 and 2006. Also, according to our predictions, the pathogen covered limited distances in the 2004-2007 period compared to the early stages of the epidemic. This quasi stasis is confirmed by the largely similar distributions of yearly incidences. Hence, here again, our approach manages to capture changes in the spatial dynamics of the pandemic that are central in the context of pathogen surveillance.

**Figure 5:**
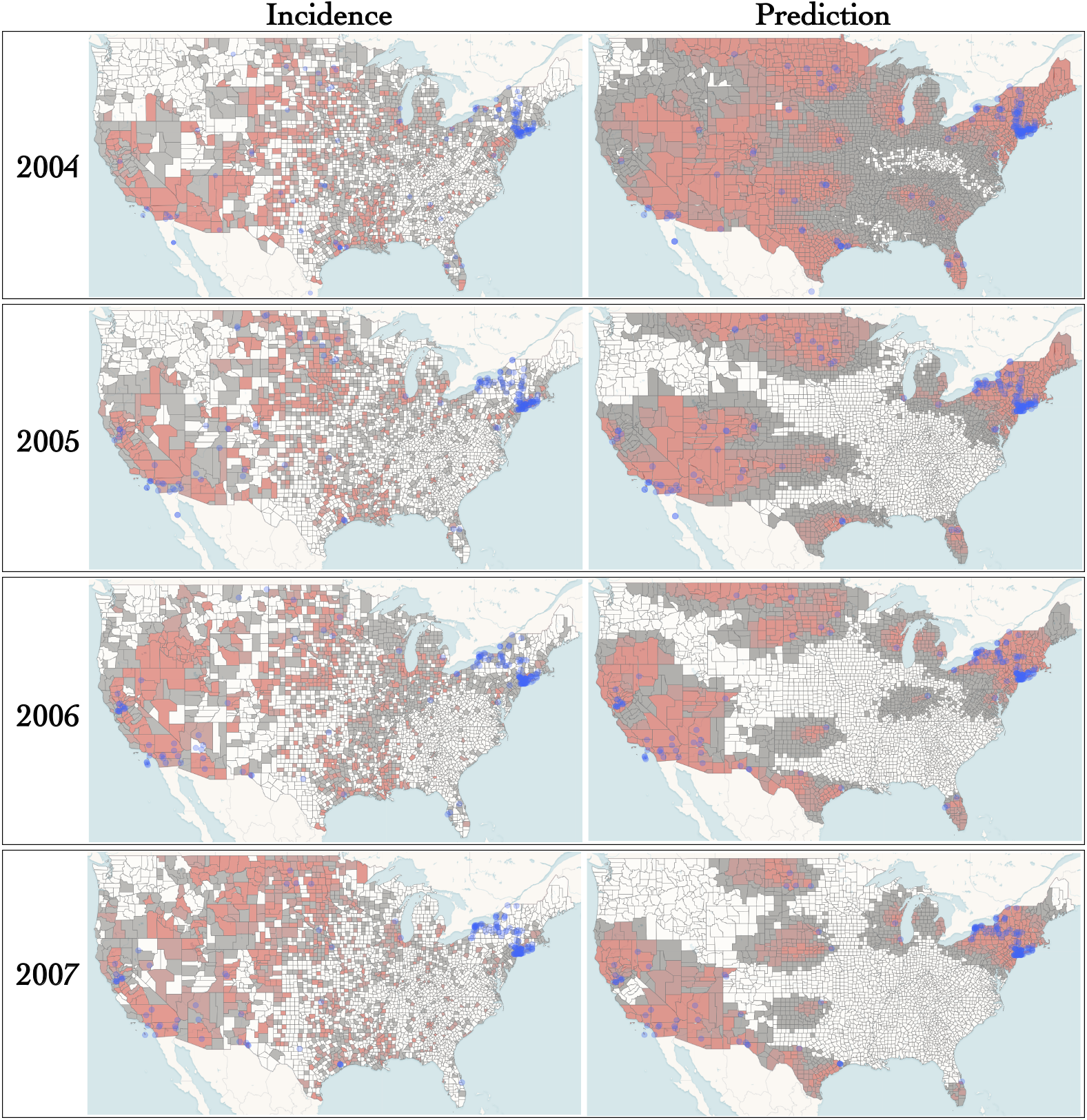
Incidence and predicted occurrence of WNV in an endemic regime (model for prediction: IBM). See caption of Fig. 4.

## Discussion

The present study addresses shortcomings in the estimation of the velocity of lineages using popular models in phylogeography. These approaches rest on the probabilistic modeling of the coordinates of lineages along their phylogeny. Yet, the central concept of instantaneous speed does not exist under the most popular Relaxed Random Walk (RRW) model. As a consequence, measuring speed as a ratio between a displacement and the corresponding elapsed time leads to difficulties. In order to circumvent these limitations, we introduce Phylogenetic Integrated Velocity (PIV) models. The originality of this family of models lies in their modeling of the velocity of evolving lineages instead of their coordinates. This approach enables a proper definition of instantaneous speed, which can be inferred anywhere along the tree, including at its tips.

Data sets were simulated under two models of spatial evolution that are distinct from that underlying the PIV and RRW approaches. Results show that speed estimates obtained with PIV models are generally more accurate than those deriving from the RRW approach, especially in cases where the pace of dispersal is high. Also, unlike RRW, PIV models produce velocity vector estimates at each node of the tree. We assessed the accuracy of these estimates at tip nodes in the IBM case, and found that the velocity vectors were well estimated, with highest posterior density intervals having good coverage (see SI, section I). Yet, PIV models are less flexible than RRW in their description of the movement of lineages during the course of evolution. In particular, sudden changes in the direction of dispersal are not well accounted for by PIV models. These changes would indeed require “breaks” in the trajectory of velocities, which the underlying Gaussian processes do not allow. However, our analysis of West Nile virus data in the U.S.A. indicates that the movements of lineages display here enough inertia so that rapid changes in the spatial trajectories are seldom observed. Cross-validation suggests in fact that the two PIV models tested here provide a fit to the data similar to that obtained with the RRW. Moreover, the analytical expressions of the variance of coordinates under the IBM model grows with time in a superlinear manner, thereby allowing large displacements in short amounts of time.

Estimates of tip velocities can serve as a basis to model future dispersal events. Here, we evaluate the accuracy of predicted movements through the analysis of subsets of a large data set of West Nile virus geo-referenced sequences and county-level yearly incidence data in the U.S.A. Our predictions focus on deciding whether the pathogen will occupy (or be absent from) a given county at a given time interval in the future, i.e., a modest, yet challenging and critical endeavor compared to predicting future incidence. The proposed approach accurately predicted the arrival of the virus along the west coast of the USA in 2002 from the analysis of data collected before the end of December 2001. Furthermore, the predictions clearly point to a change of dispersal dynamics around 2004-2005 with a transition from an expansion phase to an endemic regime whereby rapid east-to-west dispersal events are replaced with short-distance migrations. While the proposed predictions have limited accuracy in the early stages of the pandemic where data is scarce and sampling likely to be biased, the PIV models successfully anticipate dispersal events in many instances. Altogether, our results indicate that the predictive phylogeography approach put forward in the present study could indeed serve a useful purpose in real time forecasting of the spread of an epidemic. Future work could aim at incorporating data on the ecological suitability of the investigated areas in order to improve predictions, in a manner similar to that used in “landscape phylogeography” (Dellicour *et al*., 2018a).

In addition to prediction, the PIV models are also expected to prove useful in many cases where the RRW model has been applied to quantify and compare dispersal velocity. These applications range from animal and human viruses to plant viruses. For instance, lower rates of dengue virus dispersal in urban as opposed to rural settings has implicated a major role for mosquito-mediated dispersal (Raghwani *et al*., 2011). Also, dispersal velocity has often been estimated for rabies lineages with dogs as the main host species, resulting in hypotheses of their spread being impacted by human activities (Talbi *et al*., 2010). More recently, a slow dispersal has been estimated for Lassa virus in its rodent reservoir, which could in part explain the restricted distributions of the virus (Klitting *et al*., 2022). Finally, increasing dispersal rates of the rice yellow mottle virus in Africa has led to the suggestion that intensification of rice cultivation could have enhanced the spread of that virus (Rakotomalala *et al*., 2019). Applications of the PIV models could increase the credibility of these and many more hypotheses of viral spread.

The proposed new models and predictions have limitations however. In a manner similar to that of the classical RRW framework, PIV models assume that (i) the geographical position does not impact the fitness or the molecular evolution of the pathogen, (ii) all the lineages are independent from one another, excluding any competition effect and (iii) the geographical spread of the pathogen is independent from its current position. While limiting, these assumptions permit efficient computations and provided a sound methodological framework for important phylodynamics studies (see e.g. (Baele *et al*., 2018) for a review). In some specific contexts such as discrete phylogeography, some of these assumptions were relaxed, see e.g. (FitzJohn, 2010; Müller *et al*., 2017) and references therein for (i), (Drury *et al*., 2016; Manceau *et al*., 2017; Bartoszek *et al*., 2017) for (ii), and (Lemey *et al*., 2014) for (iii). Similar extensions to the PIV framework proposed here should be considered. In particular, models of animal movement also rely on integrated processes, with an additional potential function that links the dynamics of velocity evolution of an individual to its position at each point in time (Preisler *et al*., 2013; Russell *et al*., 2018). Such a potential function could be extended to include prior knowledge on the environmental layers impacting the spread of pathogens, including natural barriers such as coastline, or could be used to test the impact of specific environmental variables on the dispersion (Dellicour *et al*., 2020b). However, the pruning algorithm used here (see “Material and Methods”) would not apply to these kinds of models, which are thus likely to be highly computationally intensive.

Furthermore, sampling is likely to impact the results in case it is driven by practical aspects (e.g., the distribution of genomic surveillance facilities is not uniform throughout the habitat) and does not reflect the underlying spatial distribution of the population under scrutiny (Kalkauskas *et al*., 2021). Recent work (Guindon and De Maio, 2021) shows how different sampling strategies can be incorporated in the RRW model. A similar framework could apply to PIV models and mitigate the impact of sampling. Additionally, when available, incidence data conveys information about the demographic dynamics of an epidemic. Hence, increased accuracy of the predictions may be achievable through the incorporation of past incidence data in the new models presented in this work.

## Material and Methods

### Likelihood calculation and Bayesian inference

Let **X**^*^ and **X** correspond to random variables denoting the vectors of positions at the tips and the internal nodes respectively. **x**^*^ = *{x*_1_, …, *x*_*n*_*}* and **x** = *{x*_*n*+1_, …, *x*_2*n*−1_*}* are realizations of the corresponding random variables, where *n* is the number of tips and 2*n* − 1 is the index of the root node. **Y**^*^ and **Y** are the vectors of velocities at tip and ancestral nodes respectively. Here, we describe two different approaches for the Bayesian inference of PIV model parameters.

#### Data augmentation: sampling velocities

The first method, implemented in PhyREX (Guindon and De Maio, 2021), relies on data augmentation. It starts with the computation of *p*(**x**^*^, **x, y**^*^, **y**), i.e., the joint density of all (i.e., ancestral and tip) locations and velocities. This density is also conditioned on the phylogeny, i.e., a rooted tree topology with node heights, which is not included in the formula below for the sake of conciseness. Given the locations and velocities at all nodes in the tree, the evolutionary process taking place along every branch is independent from that happening along the other edges. The likelihood is then evaluated as follows:

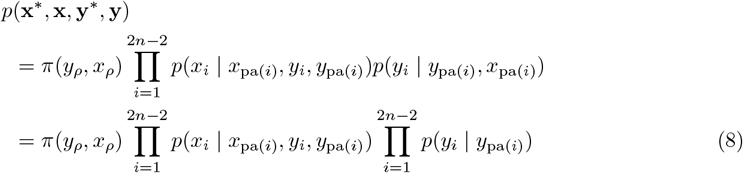

where the subscript pa(*i*) corresponds to the direct parent of node *i*. Also, *π*(*y*_*ρ*_, *x*_*ρ*_) is the velocity and location density at the root node. In the present work, we use a normal density for the corresponding distribution. Since (*X*_*i*_ | *x*_pa(*i*)_, *y*_*i*_, *y*_pa(*i*)_) is normally distributed, we can use the pruning algorithm as described in (Pybus *et al*., 2012) to integrate over **X**, giving the following likelihood:

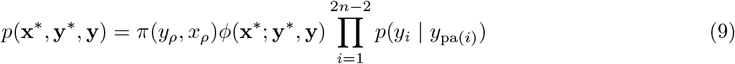

where *ϕ*(**x**^*^; **y**^*^, **y**) is obtained through a post-order tree traversal, assuming the movements along both spatial axes are independent from one another and using the means and variances for either the IBM or the IOU model (see SI, section B). The calculation just described relies on augmented data since velocities at all nodes in the tree are considered as known. Uncertainty around these latent variables is non-negligible. Samples from the joint posterior distribution of all model parameters, including ancestral and contemporaneous velocities, were obtained through Markov Chain Monte Carlo integration.

#### Direct likelihood computation with the pruning algorithm

The second method for evaluating the likelihood of PIV models is implemented in BEAST (Suchard *et al*., 2018). It relies on the direct computation of *p*(**x**^*^), the likelihood of the observed positions at the tips conditionally on the tree. It uses the fact that the stochastic process **Z**(*t*) = (**Y**(*t*), **X**(*t*)) that describes the joint evolution of both the velocity and position is a multivariate Markov process, that can be framed as linear Gaussian as in (Mitov *et al*., 2020; Bastide *et al*., 2021). Indeed, as shown in SI (section C), for any node *i* with parent pa(*i*), the joint velocity-position vector **Z**_*i*_ can be written conditionally on **Z**_pa(*i*)_ the vector at the parent node pa(*i*), as: **Z**_*i*_ = **q**_*i*_**Z**_pa(*i*)_ + **r**_*i*_ + *ϵ*_*i*_, with *ϵ*_*i*_ a Gaussian random variable with variance **σ**_*i*_ that is independent from **Z**_pa(*i*)_, and **σ**_*i*_, **q**_*i*_ and **r**_*i*_, two matrices and a vector of dimension 4 that only depend on the tree and the parameters of the PIV process considered. In this approach, all the velocities at the tips are considered as missing: we only observe the last two entries of vector **Z** corresponding to the position, but the velocities are unknown. In (Bastide *et al*., 2021), a general pruning algorithm is described to deal with this kind of process (with missing values), that provides not only the likelihood (one post-order traversal) but also the conditional distribution of non-observed traits conditioned on observed traits at the tips (one additional pre-order traversal). This algorithm hence readily gives the posterior distribution of velocities without the need to sample from them using MCMC. Moreover, it does not need to assume that movements along the spatial axes are independent from one another.

#### Phylogeographic Bayesian inference

In both approaches, standard operators were used to update the topology of the phylogenetic tree, the node ages along with the parameters of a HKY (Hasegawa *et al*., 1985a) nucleotide substitution model. The diffusion parameters of the Brownian process were also updated using standard Metropolis-Hastings steps. Most results in this study were derived with PhyREX even though BEAST outperformed PhyREX in terms of speed of parameter inference (see SI, section E). The two independent implementations of Bayesian samplers under the same models provide a robust validation of most results presented in this study.

### Simulations

#### Spatial Lambda-Fleming-Viot model

Genealogies and the accompanying spatial coordinates were first generated according to the”individual-based” spatial Lambda-Fleming-Viot (SLFV) model (Etheridge, 2008; Barton *et al*., 2010). In this model, individuals give birth to descendants which locations are normally distributed. Death events are also governed by the same kernel so that the spatial density of the population is constant, on average, during the course of evolution. The normal kernel is truncated, allowing the SLFV model to accommodate habitats of finite size, as opposed to most continuous phylogeographic models. We selected the SLFV model as it describes the evolution of a population of related individuals along a spatial continuum as opposed to discrete demes. It is not subject to the shortcomings that hinder other popular spatial population genetics models such as sampling inconsistency (Barton *et al*., 2013) or Felsenstein’s infamous “pain in the torus” (Felsenstein, 1975). Finally and most importantly, because lineages’ coordinates evolve here according to a jump process, the exact spatial coordinates of each lineage at each point in time can be monitored. This information may then serve as a basis to evaluate the total distance covered by all lineages in the genealogy. The ratio of this distance by the corresponding elapsed time gives an (average) speed that genuinely reflects the dispersal ability of the organisms under scrutiny.

50 individuals were sampled on a 10 by 10 square defining the habitat of the corresponding population. The rate of events where lineages die and/or give birth to descendants (the so-called REX events in (Guindon *et al*., 2016)) was set to 10^3^ events per unit of time per unit area and the variance of the normal density that defines the radius parameter in the SLFV model was chosen uniformly at random in [0.1, 0.3]. These parameter values are such that lineage jumps are short and frequent, thereby mimicking the behavior of a Brownian process (Wirtz and Guindon, 2023).

#### Ebola-like simulations

Here, we used the agent-based spatially explicit simulator implemented in the R package `nosoi` (Lequime *et al*., 2020). Parameters were chosen so as to mimic the Ebola epidemic in West Africa over a time period of 365 days, starting from a single infected host in Guéckédou (Guinea). `nosoi` is a discrete time, continuous space simulator that explicitly models within-host dynamics and between-host transmissions. It can exploit a geographic raster to simulate a full transmission tree where the geographic position of each infected host is tracked at all time. We simulated datasets using the same parameters as in (Lequime *et al*., 2020) which are informed by the literature describing human infections by Ebola. Spatial demographic data from WorldPop (www.worldpop.org) was also taken into account for these simulations.

Each host had a probability of 20% to move every day. These migrations were governed by a bivariate Gaussian distribution centered at the location of the lineage under scrutiny, with diagonal covariance matrix and equal standard deviations for longitude and latitude. The standard deviation was set constant for each simulation, and drawn from a log-normal distribution with mean and standard deviation equal to approximately 15 km in each direction. We used a raster of the entire West Africa, ensuring that no epidemic reached the border of the map within the time frame of the simulation.

As previously, we sampled 50 infected individuals randomly from the transmission tree, and extracted the sampled genealogy as well as the realized speed, that exploits the simulated position at each time of the chain. Note that the genealogies produced by these simulations are sampled through time and not ultrametric, making the estimation of speed easier.

#### Sequence simulation

In both simulation settings, edges in the obtained genealogy were rescaled so that the average length of an edge after scaling was 0.05 nucleotide substitutions per site. Nucleotide sequences were then generated under a strict clock model according to the HKY model of evolution (Hasegawa *et al*., 1985a) with transition/transversion ratio set to 4.0. 100 genealogies along with the corresponding spatial coordinates and homologous nucleotide sequences were generated this way for the SLFV and Ebola simulations.

#### Statistical inference

Each simulated data set was processed using the RRW, IBM and IOU models with independent coordinates. When considering their spatial components only, these models have have 3, 2 and 6 parameters respectively. The RRW model used a log-normal distribution of branch-specific dispersal rates, which is the standard parametrization for that model. The nucleotide substitution rate was set to its simulated value by taking the ratio of the tree length as expressed in molecular and calendar units. The tree-generating process was assumed to be Kingman’s coalescent (Kingman, 1982) with constant effective population size and a flat (improper) prior distribution on that parameter. Although sequences evolved according to a strict clock model, we used an uncorrelated relaxed clock model (Drummond *et al*., 2006) with a log-normal distribution of edge-specific substitution rate multipliers. An exponential prior with rate set to 100 was used for the variance of this log-normal density.

For each data set, the true average speed was taken as the actual euclidean (SLFV) and great-circle (Ebola) distance covered by every lineage divided by the tree length in calendar time unit. For RRW, distances between the (observed or estimated) coordinates at each end of every branch in the tree were used to derive the dispersal rate through the “weighted lineage dispersal velocity” statistic (Dellicour *et al*., 2016). The posterior median of that statistic was used as our speed estimate. For PIV models, speed at the tree-level was obtained by averaging the speed estimated at each node, the latter deriving from the corresponding velocities. Here again, we obtained the posterior distribution of the tree-level speed and use the median as our estimate. Note that none of the processes used for inference is the “true” process used for simulation, but simplified versions of it.

### Predictive phylogeography

The PIV models provide an adequate framework to estimate velocities at the tips of the inferred phylogenies. It thus makes sense to apply them to predicting dispersal patterns. Here, we designed a prediction technique which goal is to assess whether the organism under scrutiny may be found in a given region at a given point in time after the most recent sample was collected. Our approach utilizes the posterior distribution of the velocities estimates at each tip of the phylogeny to build a predictor. The latter is obtained by linear extrapolation of the estimated velocity at each tip in the tree that assumes a constant speed of lineages after their sampling. Survival of these linear trajectories is taken into account so that older samples are less likely than recent ones to survive to a given time point in the future. This approach therefore puts more weight on recent samples to predict dispersal patterns (see SI, section F). Incidence data used for comparison was extracted from https://www.cdc.gov/west-nile-virus/data-maps/historic-data.html.

## Data and code availability

The data and code to reproduce all analyses and figures displayed in this study are available at https://github.com/pbastide/integrated_phylogenetic_models. The PhyREX and BEAST programs are open source and freely available from https://github.com/stephaneguindon/phyml and https://github.com/beast-dev/beast-mcmc respectively.

## Acknowledgements

SG thanks the Institut Français de Bioinformatique for computational resources. The research leading to these results has received funding from the European Research Council under the European Union’s Horizon 2020 research and innovation programme (grant agreement no. 725422 - ReservoirDOCS) and the US National Institutes of Health (R01 AI153044, F31 AI154824). PR’s internship at the University of Montpellier was founded by the I-SITE MUSE through the Key Initiative “Data and Life Sciences”. PB thanks Pierre Gloaguen for useful discussions on the integrated models. SD acknowledges support from the Fonds National de la Recherche Scientifique (F.R.S.-FNRS, Belgium; grant n°F.4515.22) and the Research Foundation — Flanders (Fonds voor Wetenschappelijk Onderzoek — Vlaanderen, FWO, Belgium; grant n°G098321N). PL acknowledges support by the Research Foundation – Flanders (Fonds voor Wetenschappelijk Onderzoek – Vlaanderen, G051322N and G005323N).

## A Distribution of coordinates for the IOU process conditioned on the initial velocity

The distribution is normal with expectation and variance given below:

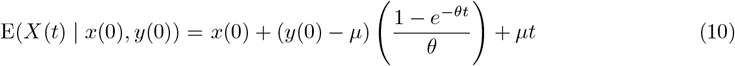

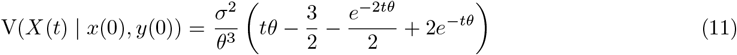

On expectation, the coordinates thus grow linearly with time, with a rate given by the trend *μ* of the underlying OU, when the process has reached its equilibrium, which happens when *e*^−*θt*^ ≃ 0. The variance of *X*(*t*) grows exponentially for values of *θt* smaller than ∼ 2 and linearly beyond that. Note that when *θ* goes to 0, the variance converges to *σ*^2^*t*^3^*/*3, so that the IOU converges to an IBM.

## B Distributions of velocity and coordinates for PIV models conditioned on the initial and final velocities

### Velocities

We consider the situation where velocities at the start and the end of a branch of length *t, y*(0) and *y*(*t*), are given, and aim at deriving the distribution of the location at the end of the branch, *X*(*t*), given the location at the start, *x*(0). Characterizing this distribution is required in the calculation of the likelihood of the PIV models in the PhyREX approach (see Section “Likelihood calculation and Bayesian inference” in the “Material and Methods” of the main text).

In case the velocity evolves according to a Brownian bridge starting at *y*(0) and stopping at *y*(*t*), the corresponding process is defined as follows:

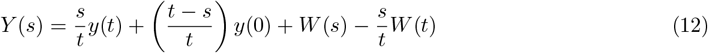

where *W* denotes the Wiener process (with *W* (0) = 0). The expected value and variance of velocity at time *s* (with 0 ≤ *s* ≤ *t*) are thus as given below:

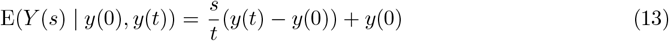

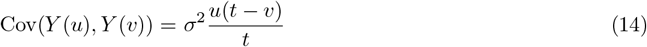

with *u* ≤ *v* ≤ *t* and *s* ≤ *t*.

When the velocity follows a OU bridge, Lemma 1 in (Papiez? and Sandison, 1990) shows that:

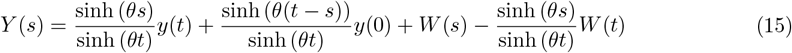

where *W* is the non-tied OU process with *W* (0) = 0. The distribution of velocity is thus normal with expectation and covariance as follows:

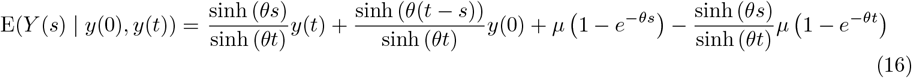

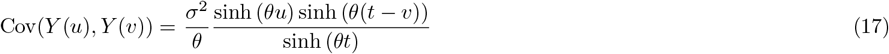

with *u* ≤ *v* ≤ *t* and *s* ≤ *t*.

### Spatial coordinates

We use the mean and covariance of the constrained velocity derived in the previous section in order to derive that of the spatial coordinates at the end of an edge of length *t*, given the coordinates at the start of that branch along with the velocity at both extremities of the same edge. We thus focus on 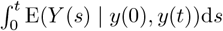 and 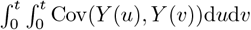 and obtain the following expressions:

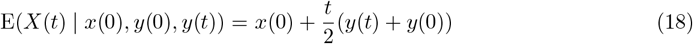

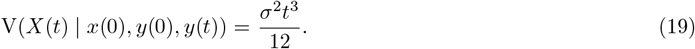

The expectation of *X*(*t*) therefore grows linearly with *t* at a pace determined by the velocity averaged over the two nodes at the extremity of the branch under scrutiny.

Using an approach equivalent to that applied to the IBM process, the expectation and variance of the coordinates given the velocities at both extremities of an edge are given below for the IOU model:

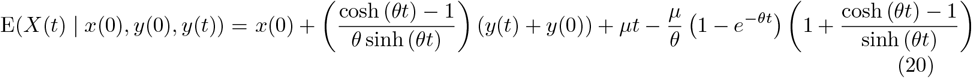

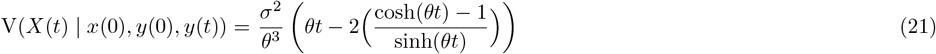

When *θ* ≪ 1, i.e., the pace to reach the equilibrium is slow with respect to *t*, we have (1 −*e*^−*θt*^)*/θ* → *t* and (cosh(*θt*) − 1)*/* sinh(*θt*) ∼ *θt/*2 so that E(*X*(*t*) | *x*(0), *y*(0), *y*(*t*)) ≃ *x*(0) + (*t/*2)(*y*(*t*) + *y*(0)). In addition, developing the variance term up to order 4 at the numerator and order 3 at the denominator, we get that (cosh(*θt*) − 1)*/* sinh(*θt*) ∼ (*θt*)*/*2 − (*θt*)^3^*/*24, so that V(*X*(*t*) | *x*(0), *y*(0), *y*(*t*)) ≃ *σ*^2^*t*^3^*/*12. As expected, when *θ* goes to 0, the IOU and the IBM processes thus behave similarly. Also, note that the function *f* : *x 1*↦ *x* − 2(cosh(*x*) − 1)*/* sinh(*x*) converges to 0 when *x* ↦ 0^+^ and the derivative of *f* with respect to *x* is non negative, so that *f* is non negative for *x* ≥ 0, and V(*X*(*t*) | *x*(0), *y*(0), *y*(*t*)) as expressed above is non negative for all values of *θt* ≥ 0. If *θ* is large, then E(*X*(*t*) | *x*(0), *y*(0), *y*(*t*)) ≃ *x*(0) + ((*y*(*t*) − *μ*) + (*y*(0) − *μ*))*/θ* + *μt*) ≃ *x*(0) + *μt*, and 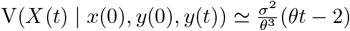.

Hence, in both regimes of *θ*, the expected value of the coordinates grows linearly with time. It is doing so in a manner that is proportional to the velocities at both extremities of the branch (if *θ* is small) or proportional to the drift term (if *θ* is large). In terms of variance, as previously, the IOU behaves similarly to the IBM when *θ* is small, and grows linearly in *t* when *θ* is large.

## C The pruning approach for the IBM and IOU processes

### Joint process

We assume here a general multivariate integrated process of dimension *p* (typically, *p* = 2 for phylogeography), and denote by **Z**(*t*) = (**Y**^*T*^ (*t*), **X**^*T*^ (*t*))^*T*^ the joint process of dimension 2*p* describing the evolution of both the velocity and position vectors. This joint process is then a linear Gaussian process, and it can be fully described by the Gaussian distribution of the trait at any node *i* given the trait at its parent pa(*i*) (Mitov *et al*., 2020; Bastide *et al*., 2021):

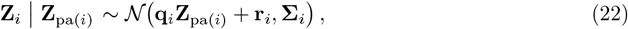

with **q**_*i*_ an actualization matrix and **σ**_*i*_ a variance matrix both of size 2*p ×* 2*p*, and **r**_*i*_ a vector or size 2*p*, that are all independent from the data, and depend only on the tree and its branch lengths. For the IBM, assuming that the velocity vector follows a BM with constant directional drift ***δ*** and variance **σ**, we get:

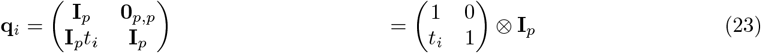

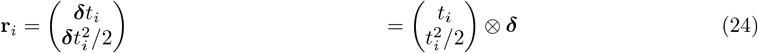

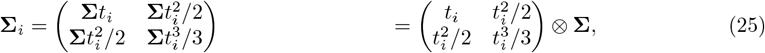

where *t*_*i*_ is the length of the branch going from pa(*i*) to *i*, and ⊗ denotes the Kronecker product. For the IOU, assuming that the velocity vector follows a OU with actualization matrix **Θ**, central vector ***μ*** and variance **σ**, we get (Cumberland and Rohde, 1977):

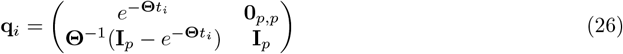

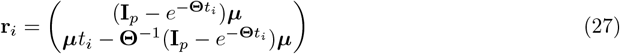

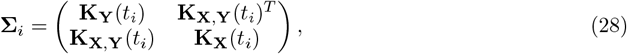

with, for any *t*, **K**_**Y**_(*t*), **K**_**X**_(*t*), and **K**_**X**,**Y**_(*t*), the variance and covariance of the velocity and position vectors at time *t*, given by:

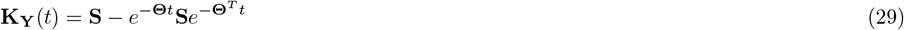

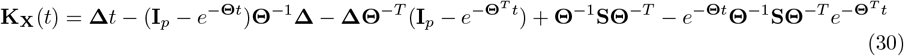

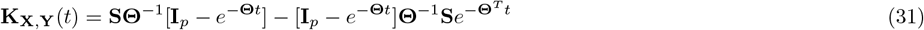

with **S** the stationary variance of the OU, and **Δ** = **Θ**^−1^**S** + **SΘ**^−*T*^.

### Pruning algorithm

As showed in (Mitov *et al*., 2020; Bastide *et al*., 2021), for a general linear Gaussian process of the form of Eq. 22 we can compute the likelihood of the traits at the tips of the tree in an efficient way by integrating over all the internal states using a pruning algorithm. This yields an algorithm in *O*(*np*^3^), that is linear in the number of tips. In the case of an integrated process, the velocity is never observed at the tips, only the spatial coordinates are. As shown in (Bastide *et al*., 2021), missing data can be accounted for in this algorithm, and because the velocity and position traits are correlated, we can still get information on the velocity, even though it is not observed. This approach is implemented in BEAST (Suchard *et al*., 2018), and allows for the Bayesian fit of the model without resorting to stochastic integration of the internal node velocities as is done in PhyREX. Note that, although we used a standard Metropolis-Hastings update for the parameters of the processes, from (Bastide *et al*., 2021) we could also get derivative with respect to the parameters of the IBM or IOU, allowing for efficient Hamiltonian Monte Carlo sampling schemes (Neal, 2011).

## D Marginal tip position distribution under the IBM and IOU processes

In the previous sections, we used the conditional distribution of a node given its parent to derive efficient pruning algorithm for the computation of the likelihood. However, as the IBM and IOU are Gaussian processes, it is also possible to directly derive the marginal distribution of the observed positions at the tip of the process. Denote by **X**_t_ the *n × p* matrix of observations at the tips of the tree, and by 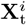 the vector of *p* observations at tip *i* 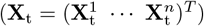.

### Marginal distribution under a BM

Recall that for a simple multivariate BM on a tree with rate variance **σ** and root position parameter **X**_*ρ*_ ∼ *𝒩* (***μ***^**X**^, **Γ**^**X**^), then we get that **X**_t_ has a matrix normal distribution, with, for any two vectors of observations at tips *i* and *j*, an expectation vector and variance covariance matrix given by:

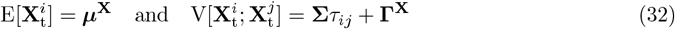

with *τ*_*ij*_ the time between the root and the most recent common ancestor of *i* and *j* (see e.g. (Felsenstein, 1973)).

### Marginal distribution under an IBM

We now assume that the pro(cess is an) IBM, with a root position parameter **X**_*ρ*_ ∼ *𝒩* (***μ***^**X**^, **Γ**^**X**^), a root velocity parameter **Y**_*ρ*_ ∼ *𝒩* **(*μ***^**Y**^, **Γ**^**Y**^), and rate variance matrix of the velocities equal to **σ**. Then the distribution of the traits at the tips is still matrix normal. More precisely, as the velocities are simply Brownian, we get that:

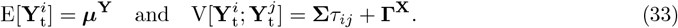

For the positions, we get:

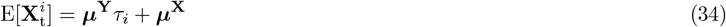

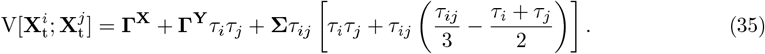

Note that, if *i* = *j*, then *τ*_*ii*_ = *τ*_*i*_, and we recover that the variance of a tip is cubic in the time of evolution. We can also get the covariances between velocities and positions:

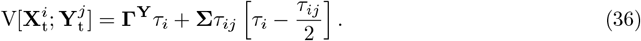

The proof of these formulas rely on the following equality:

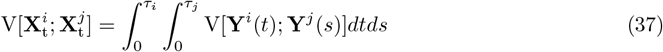

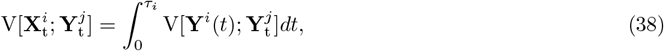

where **Y**^*i*^(*t*) denotes the value of the velocity process on lineage leading to tip *i* at time *t*, and the covariance function is equal to:

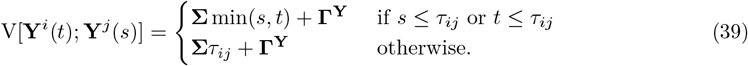

### Marginal distribution under an IOU

We now assume that the process is an IOU, with a root position parameter **X**_*ρ*_ ∼ *𝒩* (***μ***^**X**^, **Γ**^**X**^), a root velocity parameter **Y**_*ρ*_ ∼ *𝒩* (***μ***^***Y***^, ***Γ***^***Y***^), rate variance matrix of the velocities equal to **σ**, actualization matrix **Θ**, central vector ***μ*** and stationary variance **S**. Then the distribution of the traits at the tips is still matrix normal, and we get that the velocities are OU distributed as (see e.g. (Clavel *et al*., 2015)):

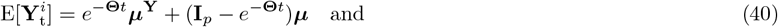

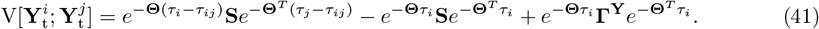

For the positions, we get:

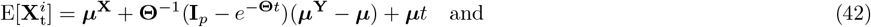

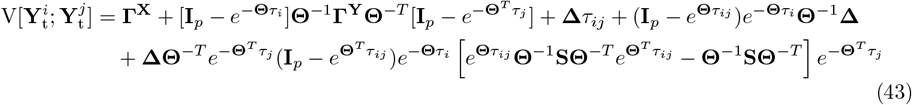

using the same notations as in section C for **Δ**.

The proof of the formulas rely on the integration of the following covariance function (see (Cumberland and Rohde, 1977)), defined, for any 0 ≤ *s* ≤ *t*, as:

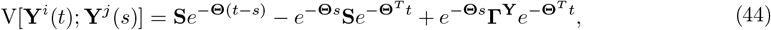

if *s* ≤ *τ*_*ij*_ or *t* ≤ *τ*_*ij*_, and: otherwise.

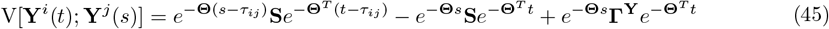

otherwise.

**Table S1:** Comparison of PhyREX and BEAST estimations of variance and root parameters for a bi-variate IBM on a fixed tree (WNV data of (Pybus *et al*., 2012) with 104 tips).

| parameter | PhyREX |  |  | BEAST |  |  |
| --- | --- | --- | --- | --- | --- | --- |
|  | ESS/s | mean | 95% HPDI | ESS/s | mean | 95% HPDI |
| $\sigma^2$ lat | 9.73 | 2.37 | ( 1.7, 3.1) | 100.15 | 2.29 | ( 1.7, 3.0) |
| $\sigma^2$ lon | 10.82 | 11.81 | ( 8.7, 15.3) | 111.90 | 11.32 | ( 8.4, 14.5) |
| root lat | 5.74 | 39.88 | ( 32.9, 47.1) | 164.66 | 40.65 | ( 33.7, 47.6) |
| root lon | 6.78 | -71.07 | ( -86.2, -55.7) | 164.11 | -74.62 | ( -90.3, -59.3) |
| root veloc lat | 17.63 | -0.13 | ( -3.4, 3.3) | 166.60 | -0.26 | ( -3.5, 3.1) |
| root veloc lon | 17.81 | -3.80 | ( -11.0, 3.7) | 161.19 | -2.32 | ( -9.6, 4.9) |

### Likelihood computation checks

The formulas above give the full distribution of the observed positions at the tips of the tree, that is matrix normal. They are therefore used to compute the likelihood of the data directly. As in the standard BM case, this direct computation involves the inversion of large matrices, and the pruning approach of section C is to be preferred, as it is linear in the number of tips (see e.g. (Mitov *et al*., 2020; Bastide *et al*., 2021)). However, these formulas can be used to test that the likelihoods obtained by BEAST and PhyREX do match with the direct “naïve” formulas. We implemented these direct formulas in `R` (R Core Team, 2024), and checked on small examples that the two software gave matching likelihoods.

## E Comparison of PhyREX and BEAST implementations

We compared two independent implementations of the IBM model in PhyREX and BEAST, with the WNV data from (Pybus *et al*., 2012) (104 tips), independent IBM models for the latitudes and longitudes, and a vague Gaussian prior centered at 0 and with variance 1000 for the root trait. We compared the two implementations first using a common fixed tree, and then starting from sequences and inferring the tree.

### Settings

For the fixed tree analysis, as PhyREX needs to sample interval velocities, we ran a longer MCMC chain for this software, with 10 million iterations sampled every 10 thousands steps, while we used a chain with only 10 thousands iterations sampled every 10 steps for BEAST. We reproduced this analysis 10 times on a 13-inch M2 2022 MacBook Pro, and took the mean times and estimates.

For the inferred tree analysis, we ran a MCMC chain with 50 million iterations sampled every 10 thousands steps for both software. Since the analyses are computationally intensive, we only ran the chain once with the same set up and compared running times. For both analyses, we used a constant population coalescent prior, an HKY substitution model (Hasegawa *et al*., 1985b), and an uncorrelated relaxed random local clock (Drummond *et al*., 2006) with a log-normal distribution of rates with a strong prior on a small standard error to speed up convergence. Tip heights were fixed at their sampling times.

We used `R` (R Core Team, 2024) to run analyses, using packages `ape` (Paradis and Schliep, 2019) and `treeio` (Wang *et al*., 2019) for tree manipulation, `tracerer` (Bilderbeek and Etienne, 2018) for reading and summarizing log files, `ggplot2` (Wickham, 2016) and `cowplot` (Wilke, 2024) for plotting results, `here` (Müller, 2020) for file manipulation, and `kableExtra` (Zhu, 2024) for extracting tables automatically for display.

### Results for the fixed tree analysis

We found that the two approaches gave very similar results for the estimation of the variance and root parameters (see Table S1). Unsurprisingly, as BEAST only needs to sample two parameters (the variance parameters), it was faster in this setting, with analyses taking around 5.2 seconds, versus 33.6 seconds with PhyREX. This led to a mean approximate 15.3 times factor increase of the BEAST versus PhyREX implementation in terms of effective sample size per seconds (see Table S1). Because velocities are sampled during the MCMC in PhyREX, but directly sampled in their posterior distribution in BEAST, their ESS in BEAST was more reliably high and less spread out than PhyREX estimations (see Fig. S1-a). Note that each PhyREX iteration was about 155.0 times faster than each BEAST iteration, as it requires less computations.

**Figure S1:**
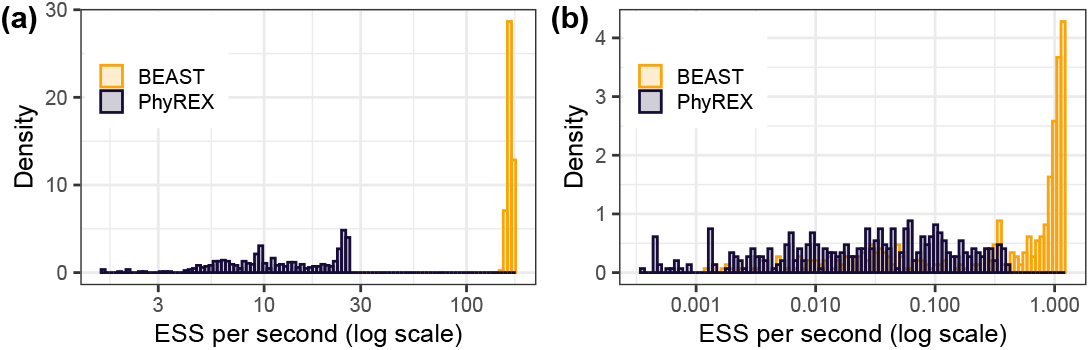
Effective sample size per seconds of all node velocities. (internal and tips) of BEAST versus PhyREX for a bi-variate IBM on a fixed tree (a) or with an inferred tree (b). WNV data of (Pybus *et al*., 2012) with 104 tips, i.e. 208 velocity estimates per software program.

**Figure S2:**
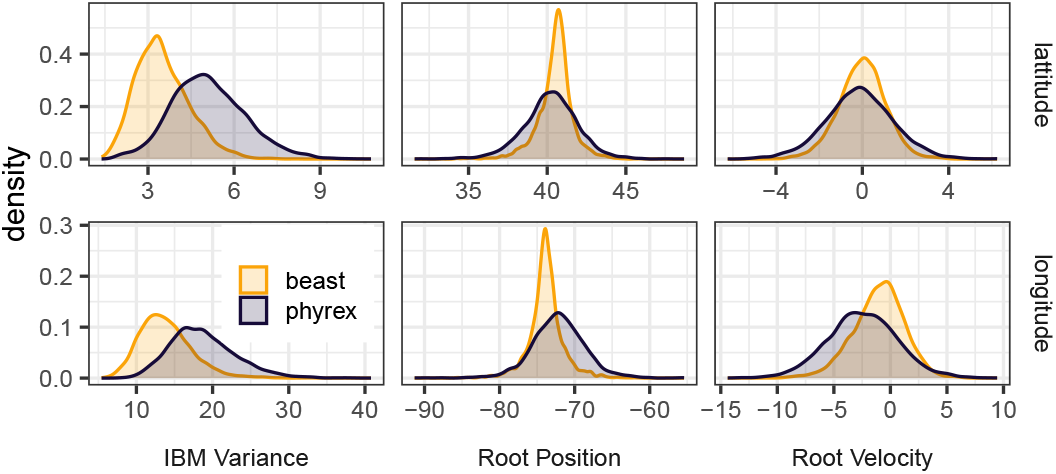
Comparison of PhyREX and BEAST estimations of variance and root parameters for a bi-variate IBM when the tree is inferred using WNV data of (Pybus *et al*., 2012) with 104 tips.

### Results for the inferred tree analysis

As inferring the tree is much more computationally intensive, these analyses took longer to run, taking, respectively, 1.05 hours for BEAST and 2.85 hours for PhyREX. PhyREX convergence was slower, taking around 10 million iterations to warm up, while BEAST reached a reasonable sampling area within less than 1 million iterations. This led to a mean approximate 13.70 times factor increase of the BEAST versus PhyREX implementation in term of effective sample size per seconds. As in the fixed tree case, ESS in BEAST was more reliably high and less spread out than PhyREX estimations (see Fig. S1-b). Because many operators are involved in the tree search, it is difficult to pinpoints the exact reasons for these different convergence behaviors, and it may not be entirely due to the different IBM implementations. Contrary to the fixed tree case, each BEAST iteration was faster than a PhyREX iteration by a factor of about 2.70. This time difference is likely due to the use of BEAGLE (Ayres *et al*., 2012) in the BEAST analyse, which allows for efficient parallelization and GPU use. The two approaches gave very similar results for the estimation of the root time and position, but gave slightly different variance parameter estimations, although with intersecting HPD intervals (see Fig. S2). Reconstructed velocities at tips were highly similar in both approaches (not shown).

## F Dispersal prediction via linear extrapolation of tip velocities

Let *α*_*i*_(*t*) design the coordinates of tip *i* at time *t* that occurs after the sample corresponding to tip *i* was observed, which is noted as *t*_*i*_ (i.e. *t* ≥ *t*_*i*_). *α*_*i*_(*t*) then corresponds to the position of lineage *i*, should it survive up to time *t*. Also, we assume that, after time *t*_*i*_, lineage *i* dies at rate *λ* so that the probability of surviving up to time *t* is exp(−*λ*(*t* − *t*_*i*_)). Finally, *𝒜* represents a particular region, e.g., a state or county. Below is the probability that one or more lineage occupies *𝒜* at time *t*, where *t* ≥ *t*_*i*_ for all tips *i* = 1, …, *n*. Let *𝒫* _*𝒜,t*_ be that probability. We have:

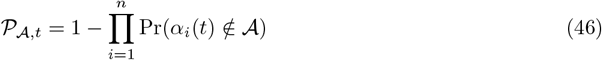

where Pr(*α*_*i*_(*t*) ∉ *𝒜*) is the probability that lineage *i* is not within *𝒜* at time *t*. We have Pr(*α*_*i*_(*t*) ∉ *𝒜*) = 1 − Pr(*α*_*i*_(*t*) ∈ *𝒜*) and Pr(*α*_*i*_(*t*) ∈ *𝒜*) is the probability that (1) lineage *i* survives up to time *t* and (2) the linear extrapolation of its position from time *t*_*i*_ to time *t* given its velocity at time *t*_*i*_, falls within *𝒜*. In practice, we are interested in the probability of occupation for a given time interval [*t, t* + *s*], which we approximate as follows:

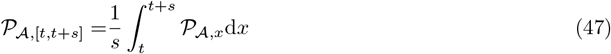

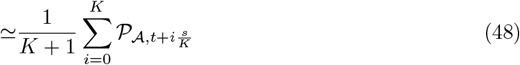

The time unit considered in this study is the year and we used *K* = 4 so that one year is split up into four parts of equal lengths. Also, we fixed the value of *λ* to 1.0 so that the probability of a lineage to survive for a period of one year is 0.37. This value of *λ* derived from the observation that the length of external branches in WNV phylogenies are generally close to 1.0. Assuming a critical birth-death model approximates the branching process here, Theorem 3 in (Mooers *et al*., 2012) states that the rate of death of lineage is given by the inverse of the average length of an external edge. Note that PIV models allow us to derive the joint distribution of the velocity and position of each tip conditionally on all other tips thanks to the pruning algorithm described in Section C. We could therefore derive the distribution of the particle starting from this tip after a time *t*, assuming that the velocity is constant. Probabilities *𝒫* _*𝒜,x*_ would then be obtained by integrating this distribution over on domain *𝒜*. Such an approach could be computationally expensive and would need to be carefully examined for possible use in future work.

## G Cross-validation of tip coordinates

In an attempt to compare the fit of the PIV and the RRW models to the WNV data sets, we assessed the ability of these two models to recover coordinates at tips where only sequence data is made available. A MCMC analysis was first performed on the full data set. Then, for each tip taken in a sequential manner, coordinates were hidden and considered as parameters of the model. The posterior distribution of the standard model parameters (including the tree topology, age of internal nodes plus the dispersal parameters of the model considered) along with that of the missing tip location, were obtained and the posterior distribution of the great circle distance between the true and estimated tip locations was recorded.

Let 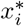 be the tip coordinates at tip *i* and 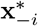 the set of coordinates observed at all tips except *i*. Also, let **s**^*^ be the set of observed sequences at the tips. *θ* is a generic parameter that encompasses all the parameters of the model excluding the tip and ancestral velocities, noted as **y**^*^ and **y** respectively. Our objective is to draw samples from the distribution of 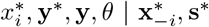. Given samples from this joint distribution, one marginalizes over **y**^*^, **y** and *θ* in order to recover the posterior distribution of interest. We have:

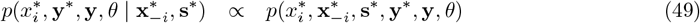

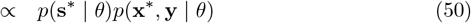

so that samples from the target distribution can be obtained by standard MCMC with *x*_*i*_ considered as a parameter of the model along **y**^*^, **y** and *θ*. As explained above, we ran a first MCMC on the full data set so as to reach the stationary distribution of the Markov chain that generates correlated samples from *p*(**y**^*^, **y**, *θ* | **x**^*^, **s**^*^). Each tip coordinates is then hidden sequentially. For each hidden tip coordinates 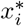, a shorter MCMC analysis is ran with 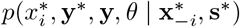 as its target distribution.

**Figure S3:**
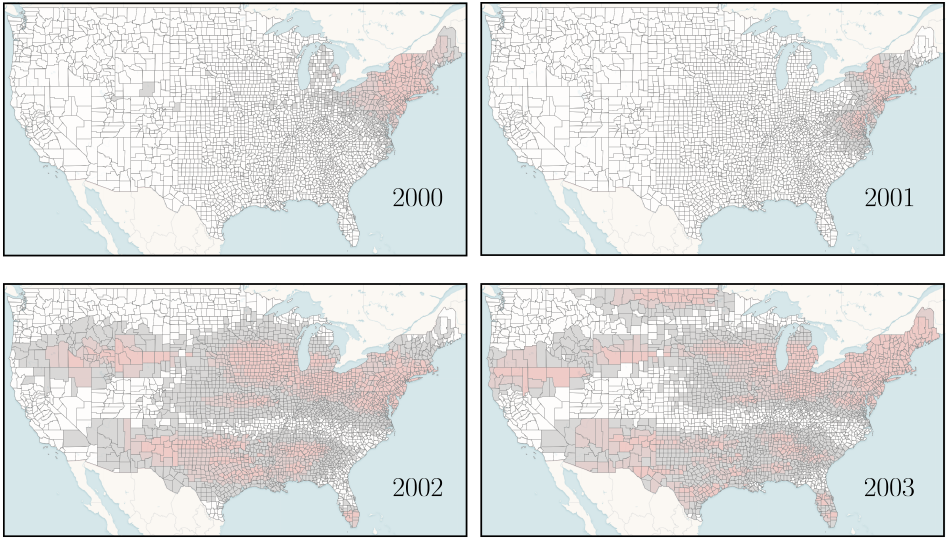
Predicted occurrence of WNV in the early phase of the epidemic (model for prediction: IBM). Here, the prior distribution for the velocity at the root had a variance set to (10^−2^, 10^−2^) (mean set to (0, 0)) as opposed to a virtually flat prior (variance set to (10^2^, 10^2^)) by default (see Fig. 4 of the main text).

Note that, thanks to the pruning algorithm described in Section C, one could directly get the distribution of left-out tips conditionally on observed ones, to carry out a cross validation relying on the expected log predictive density similar to (Hassler *et al*., 2022).

## H Prediction of WNV incidence deriving from alternative models

By default, predictions were performed using the IBM model with a flat prior on each of the two parameters making up the variance of the velocity vector. A normal distribution centered on (0, 0) and variance (10^2^, 10^2^) was used here. Figure S3 gives the predicted occurrence of WNV in the early stages of the epidemic using a more informative prior with variance vector (10^−2^, 10^−2^). While the predictions for years 2001-2003 are similar to that obtained with a flat prior, the occupied area for year 2000 is smaller when using the informative prior compared to that obtained with a non-informative one. The sensitivity to priors observed here is a likely consequence of the lack of signal conveyed by the limited amount of data (only seven sequences with coordinates are available for this time point).

We also performed prediction analyses using the RRW model. Velocity at each tip was estimated during the MCMC analysis as follows: (1) ancestral location were sampled from their joint posterior density; (2) great-circle distances between each tip location and that sampled for its direct ancestor were evaluated; (3) the obtained distance was divided by the time elapsed along the corresponding (external) edge. Predictions were then made using the same approach as that used with the IBM model (see SI, section F). Figures S4 and S5 show the incidence from the CDC data (see main text) and the corresponding predictions using the RRW model. We note that the predictions of the RRW are much more spread out than the ones of the IBM (see main text and Figure S3), which is consistent with the fact that the RRW can allow for jumps in the process, i.e. large dispersion events in small time scales, so that, according to this model, a spread far away from the origin is not unlikely, even in the early phases of the epidemics.

We compared the IBM and RRW models by evaluating their sensitivity (true positive rate) and specificity (true negative rate) corresponding to the predicted occurrence of the virus in each county. More specifically, for each year in the 2000-2007 time period, a county was said to be predicted as “infected” as soon as at least one lineage was predicted in this county. This county was then labeled as a “true positive” if the corresponding incidence from the CDC data (see main text) showed at least one case. Figure S6 shows that the RRW has a greater sensitivity, but a lower sensitivity. This is consistent with the spatial patterns observed on Fig. S5 and S4, that showed a wider dispersion for the RRW.

Using the empirical cumulative distribution function, we transformed the predicted count data of both methods into probabilities. These probabilities were then used to predict the occurrence of the virus in each county. This allowed us to compute Receiver Operating Characteristic (ROC) curve for both predictor for each year, using the R package `ROCR` (Sing *et al*., 2005). Results in Fig. S7 show that the IBM predictor is generally further from the diagonal than the RRW predictor, indicating a better overall performance of the IBM. We also note that the predictor are more accurate for the early stages of the epidemics, when the spatial distribution of the virus is limited.

**Figure S4:**
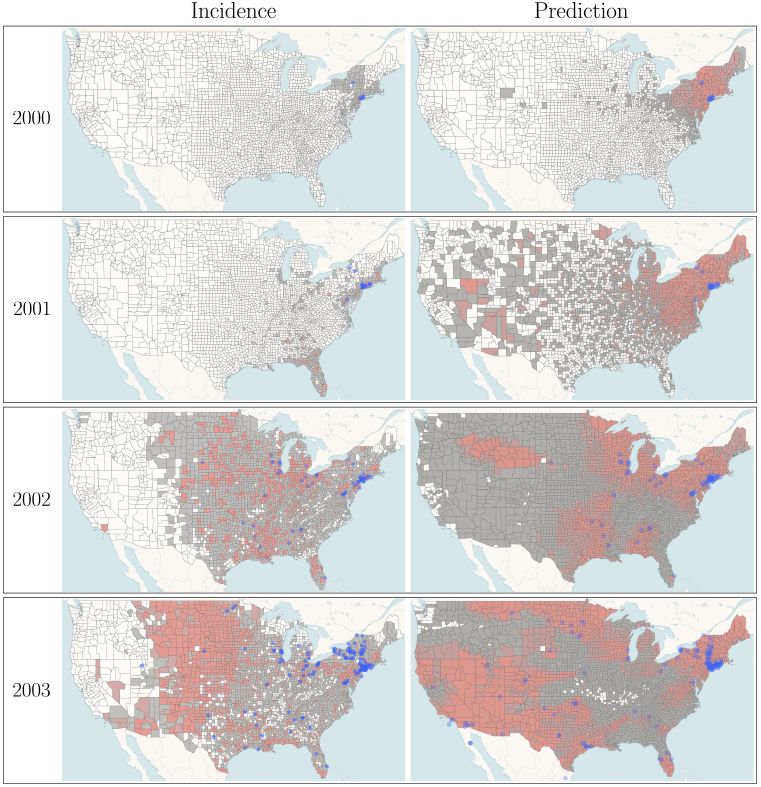
Incidence and predicted occurrence of WNV in the early phase of the epidemic (model for prediction: RRW). Purple dots correspond to sampled locations. Incidence data (left) for each year and each county was obtained from the CDC. For year *Y*, predicted occurrence of the WNV (right) was inferred using data collected earlier than the end of December of year *Y* − 1. The maps were generated with EvoLaps2 (Chevenet *et al*., 2024)

**Figure S5:**
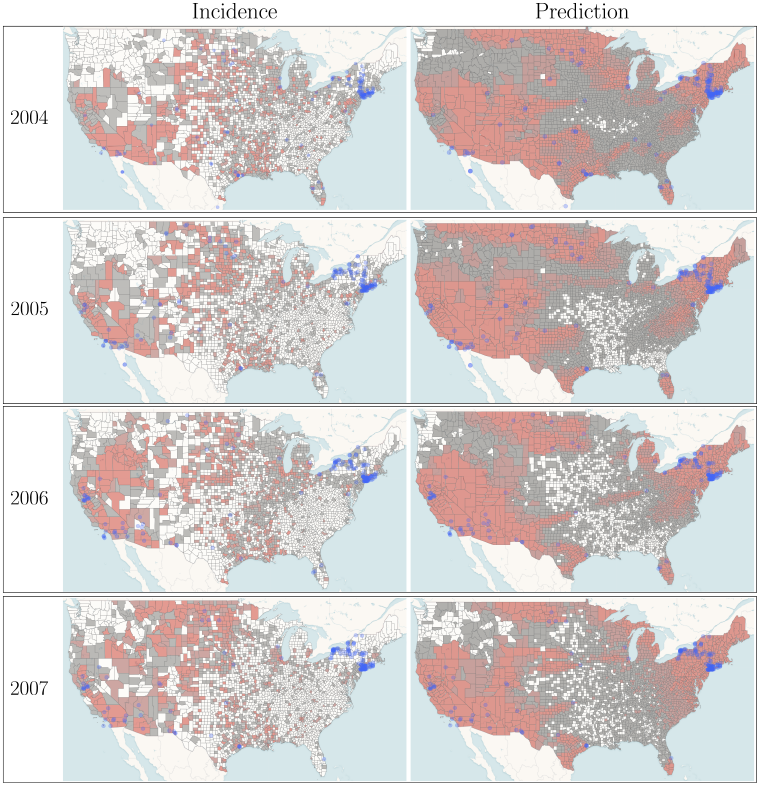
Incidence and predicted occurrence of WNV in an endemic regime (model for prediction: RRW). See caption of Figure S4.

**Figure S6:**
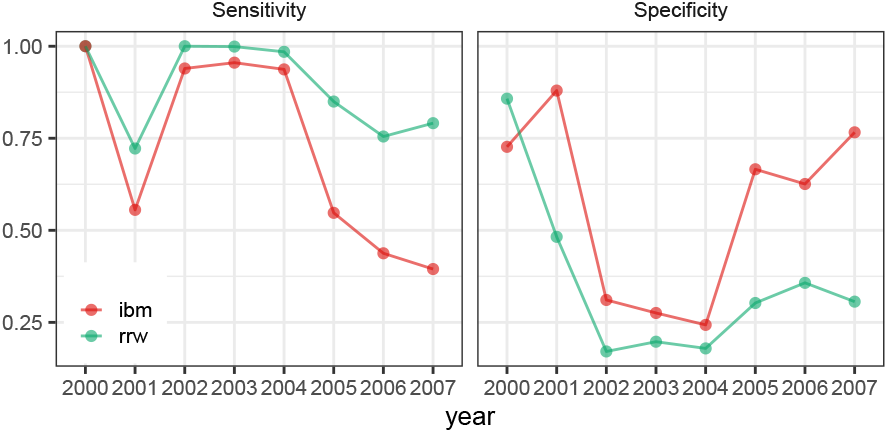
Sensitivity and specificity of the predicted county level occurrences by the IBM and RRW models.

**Figure S7:**
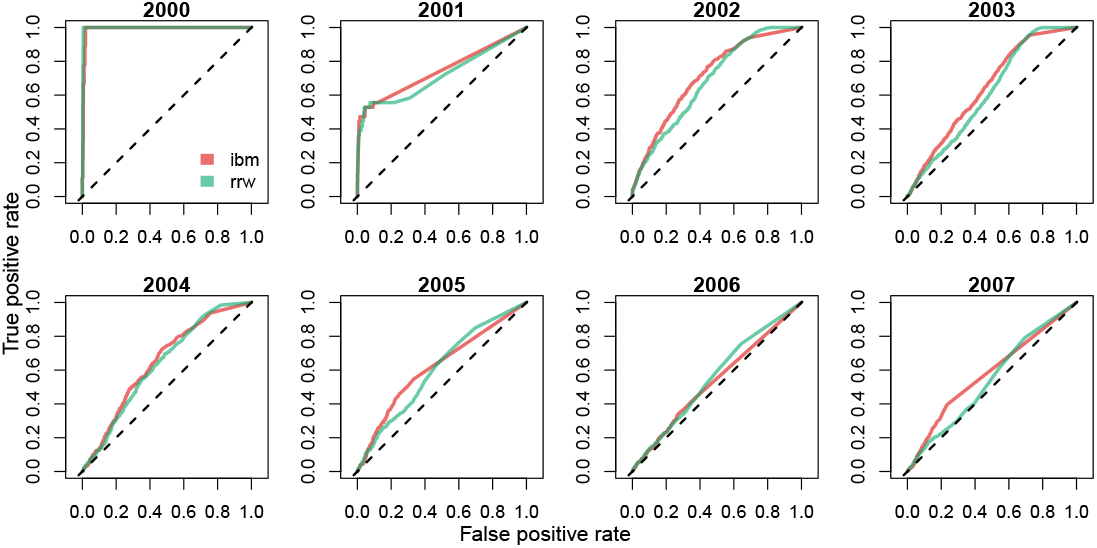
ROC curves for the predicted county level occurrences by the IBM and RRW models.

## I Tip velocity estimation under the IBM model

We performed a simulation study using the IBM both for simulation and inference in order to check that our model and implementation produced correct results when presented with datasets that matched its assumptions.

We took a fixed and dated WNV tree inferred from previous analyses, and simulated 100 datasets using an IBM with independent movements on the latitude and longitude axes, with variances respectively set to 0.1 and 1. The root location was set to the New-York region (latitude 40.65°, longitude -74.33°) with root velocity vector (−0.24, -2.48) degrees per year.

We then used the BEAST implementation of the IBM, using the true fixed tree, but inferring all other parameters from the data. For each dataset, we ran an MCMC chain for 50 000 iterations, log every 100, with a standard log-transformed random walk operator on the variance parameters, and vague half-t priors. We then extracted estimates and 95% highest posterior density intervals (HPDI) for variance parameters, root position and velocity, and all tip velocity vectors.

Fig. S8 shows that the true variance parameters and root position and velocity are correctly recovered, with unbiased estimates, as expected. Further, Fig. S9 shows that the 104 tip velocity vectors are also correctly estimated, with unbiased estimates, and HPDI reaching coverages close to their nominal values.

In this simple setting with a fixed tree and an IBM used for both simulation and inference, this experiment shows that our implementation can recover the correct velocity dynamic of the epidemic over the different regions of the tree. Further investigations, using other velocity-explicit simulation models, could be the focus of future work which goal would be to asses the robustness of PIV models and its ability to recover specific propagation dynamics.

**Figure S8:**
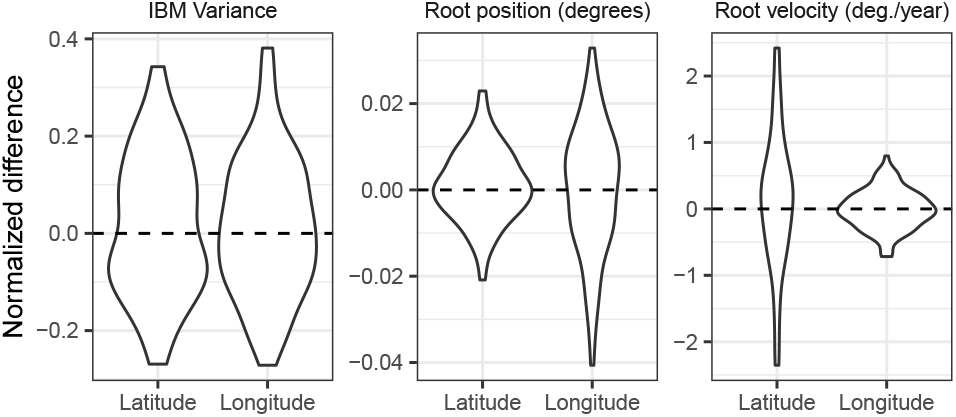
Variance and root parameter IBM estimate. Difference between the estimated and true value, normalized by the true value, for the IBM variance parameter (first panel) and the root position and velocity vectors (in degrees, second and last panels). Violin plot over 100 replicates. Data was simulated using an IBM on the WNV tree with 104 tips. BEAST was used for inference, using the true fixed tree, and an IBM model.

**Figure S9:**
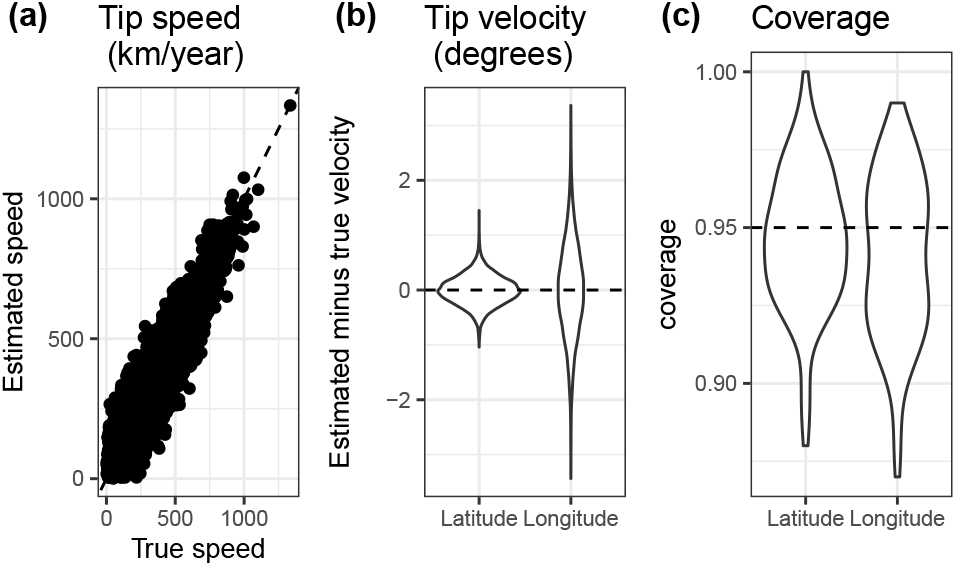
Tip speed and velocity estimates. **(a)** Estimated vs. true speed (in km/year) for the 104 tips and the 100 replicates. **(b)** Estimated minus true tip velocity vector (in degrees). **(c)** Realized coverage of the 95% highest posterior density intervals for the tip velocity vectors. Violin plot over 100 replicates and 104 tips. Data was simulated using an IBM on the WNV tree with 104 tips. BEAST was used for inference, using the true fixed tree, and an IBM model.

## References

Ayres, D., Darling, A., Zwickl, D., Beerli, P., Holder, M., Lewis, P., Huelsenbeck, J., Ronquist, F., Swofford, D., Cummings, M., et al. 2012. BEAGLE: an application programming interface and high-performance computing library for statistical phylogenetics. Systematic Biology, 61: 170–173.

Baele, G., Dellicour, S., Suchard, M., Lemey, P., and Vrancken, B. 2018. Recent advances in computational phylodynamics. Current Opinion in Virology, 31: 24–32.

Barndorff-Nielsen, O. and Shephard, N. 2003. Integrated OU processes and non-Gaussian OU-based stochastic volatility models. Scandinavian Journal of Statistics, 30: 277–295.

Barton, N., Etheridge, A., and Véber, A. 2010. A new model for evolution in a spatial continuum. Electronic Journal of Probability, 15: 162–216.

Barton, N., Etheridge, A., and Véber, A. 2013. Modelling evolution in a spatial continuum. Journal of Statistical Mechanics: Theory and Experiment, 2013: P01002.

Bartoszek, K., Glémin, S., Kaj, I., and Lascoux, M. 2017. Using the Ornstein–Uhlenbeck process to model the evolution of interacting populations. Journal of Theoretical Biology, 429: 35–45.

Bastide, P. and Didier, G. 2023. The Cauchy process on phylogenies: A tractable model for pulsed evolution. Systematic Biology, 72: 1296–1315.

Bastide, P., Ho, L. S. T., Baele, G., Lemey, P., and Suchard, M. 2021. Efficient Bayesian inference of general Gaussian models on large phylogenetic trees. The Annals of Applied Statistics, 15: 971–997.

Bilderbeek, R. and Etienne, R. 2018. babette: Beauti 2, beast2 and tracer for r. Methods in Ecology and Evolution, 9(9): 2034–2040.

Campbell, G., Marfin, A., Lanciotti, R., and Gubler, D. 2002. West Nile virus. The Lancet Infectious Diseases, 2: 519–529.

Chevenet, F., Fargette, D., Bastide, P., Vitré, T., and Guindon, S. 2024. EvoLaps 2: Advanced phylogeographic visualization. Virus Evolution, 10: vead078.

Clavel, J., Escarguel, G., and Merceron, G. 2015. mvmorph : an R package for fitting multivariate evolutionary models to morphometric data. Methods in Ecology and Evolution, 6: 1311–1319.

Cornuault, J. 2022. Bayesian analyses of comparative data with the Ornstein–Uhlenbeck model: potential pitfalls. Systematic Biology, 71: 1524–1540.

Cumberland, W. and Rohde, C. 1977. A multivariate model for growth of populations. Theoretical Population Biology, 11: 127–139.

Dellicour, S., Rose, R., Faria, N. R., Lemey, P., and Pybus, O. G. 2016. SERAPHIM: studying environmental rasters and phylogenetically informed movements. Bioinformatics, 32(20): 3204– 3206.

Dellicour, S., Rose, R., Faria, N., Vieira, L., Bourhy, H., Gilbert, M., Lemey, P., and Pybus, O. 2017. Using viral gene sequences to compare and explain the heterogeneous spatial dynamics of virus epidemics. Molecular Biology and Evolution, 34: 2563–2571.

Dellicour, S., Vrancken, B., Trovão, N., Fargette, D., and Lemey, P. 2018a. On the importance of negative controls in viral landscape phylogeography. Virus Evolution, 4: vey023.

Dellicour, S., Baele, G., Dudas, G., Faria, N., Pybus, O., Suchard, M., Rambaut, A., and Lemey, P. 2018b. Phylodynamic assessment of intervention strategies for the West African Ebola virus outbreak. Nature Communications, 9: 1–9.

Dellicour, S., Lequime, S., Vrancken, B., Gill, M., Bastide, P., Gangavarapu, K., Matteson, N., Tan, Y., Du Plessis, L., Fisher, A., et al. 2020a. Epidemiological hypothesis testing using a phylogeographic and phylodynamic framework. Nature Communications, 11: 5620.

Dellicour, S., Lemey, P., Artois, J., Lam, T., Fusaro, A., Monne, I., Cattoli, G., Kuznetsov, D., Xenarios, I., Dauphin, G., et al. 2020b. Incorporating heterogeneous sampling probabilities in continuous phylogeographic inference—application to H5N1 spread in the Mekong region. Bioinformatics, 36(7): 2098–2104.

Dellicour, S., Bastide, P., Rocu, P., Fargette, D., Hardy, O., Suchard, M., Guindon, S., and Lemey, P. 2024. How fast are viruses spreading in the wild? bioRxiv.

Drummond, A., Pybus, O., Rambaut, A., Forsberg, R., and Rodrigo, A. 2003. Measurably evolving populations. Trends in Ecology and Evolution, 18: 481–488.

Drummond, A., Ho, S., Phillips, M., and Rambaut, A. 2006. Relaxed phylogenetics and dating with confidence. PLoS Biology, 4: e88.

Drury, J., Clavel, J., Manceau, M., and Morlon, H. 2016. Estimating the effect of competition on trait evolution using maximum likelihood inference. Systematic Biology, 65: 700–710.

Etheridge, A. 2008. Drift, draft and structure: some mathematical models of evolution. Banach center publications, 80: 121–144.

Felsenstein, J. 1973. Maximum-likelihood estimation of evolutionary trees from continuous characters. The American Journal of Human Genetics, 25: 471–492.

Felsenstein, J. 1975. A pain in the torus: some difficulties with models of isolation by distance. American Naturalist, 109: 359–368.

FitzJohn, R. 2010. Quantitative traits and diversification. Systematic Biology, 59: 619–633.

Gardiner, C. 2009. Stochastic Methods. Springer Berlin, Heidelberg, 4th edition.

Gill, M., Tung H. L.S.,, Baele, G., Lemey, P., and Suchard, M. 2016. A relaxed directional random walk model for phylogenetic trait evolution. Systematic Biology, 66: 299–319.

Guindon, S. and De Maio, N. 2021. Accounting for spatial sampling patterns in Bayesian phylogeography. Proceedings of the National Academy of Sciences, 118: e2105273118.

Guindon, S., Guo, H., and Welch, D. 2016. Demographic inference under the coalescent in a spatial continuum. Theoretical Population Biology, 111: 43–50.

Hasegawa, M., Kishino, H., and Yano, T. 1985a. Dating of the Human-Ape splitting by a molecular clock of mitochondrial-DNA. Journal of Molecular Evolution, 22: 160–174.

Hasegawa, M., Kishino, H., and Yano, T. 1985b. Dating of the Human-Ape splitting by a molecular clock of mitochondrial-DNA. Journal of Molecular Evolution, 22: 160–174.

Hassler, G., Gallone, B., Aristide, L., Allen, W., Tolkoff, M., Holbrook, A., Baele, G., Lemey, P., and Suchard, M. 2022. Principled, practical, flexible, fast: A new approach to phylogenetic factor analysis. Methods in Ecology and Evolution, 13: 2181–2197.

Ho, S. and Shapiro, B. 2011. Skyline-plot methods for estimating demographic history from nucleotide sequences. Molecular Ecology Resources, 11: 423–434.

Holmes, E. 2013. What can we predict about viral evolution and emergence? Current Opinion in Virology, 3: 180–184.

Holmes, E. C., Rambaut, A., and Andersen, K. G. 2018. Pandemics: spend on surveillance, not prediction. Nature, 558(7709): 180–182.

Hooten, M. and Johnson, D. 2017. Basis function models for animal movement. Journal of the American Statistical Association, 112: 578–589.

Issaka, S., Traoré, O., Longué, R. D. S., Pinel-Galzi, A., Gill, M., Dellicour, S., Bastide, P., Guindon, S., Hébrard, E., Dugué, M.-J., et al. 2021. Rivers and landscape ecology of a plant virus, Rice yellow mottle virus along the Niger Valley. Virus Evolution, 7: veab072.

Johnson, D., London, J., Lea, M.-A., and Durban, J. 2008. Continuous-time correlated random walk model for animal telemetry data. Ecology, 89: 1208–1215.

Kalkauskas, A., Perron, U., Sun, Y., Goldman, N., Baele, G., Guindon, S., and De Maio, N. 2021. Sampling bias and model choice in continuous phylogeography: getting lost on a random walk. PLOS Computational Biology, 17(1): e1008561.

Kingman, J. 1982. The coalescent. Stochastic Processes and their Applications, 13: 235–248.

Klitting, R., Kafetzopoulou, L., Thiery, W., Dudas, G., Gryseels, S., Kotamarthi, A., Vrancken, B., Gangavarapu, K., Momoh, M., Sandi, J., et al. 2022. Predicting the evolution of the Lassa virus endemic area and population at risk over the next decades. Nature Communications, 13: 5596.

Lanciotti, R., Roehrig, J., Deubel, V., Smith, J., Parker, M., Steele, K., Crise, B., Volpe, K., Crabtree, M., Scherret, J., et al. 1999. Origin of the West Nile virus responsible for an outbreak of encephalitis in the northeastern United States. Science, 286(5448): 2333–2337.

Lemey, P., Rambaut, A., Welch, J., and Suchard, M. 2010. Phylogeography takes a relaxed random walk in continuous space and time. Molecular Biology and Evolution, 27: 1877–1885.

Lemey, P., Rambaut, A., Bedford, T., Faria, N., Bielejec, F., Baele, G., Russell, C., Smith, D., Pybus, O., Brockmann, D., and Suchard, M. 2014. Unifying viral genetics and human transportation data to predict the global transmission dynamics of Human Influenza H3N2. PLoS Pathogens, 10(2): e1003932.

Lequime, S., Bastide, P., Dellicour, S., Lemey, P., and Baele, G. 2020. nosoi: A stochastic agent-based transmission chain simulation framework in R. Methods in Ecology and Evolution, 11: 1002–1007.

Malécot, G. 1948. Mathematics of heredity. Paris: Masson et Cie.

Manceau, M., Lambert, A., and Morlon, H. 2017. A unifying comparative phylogenetic framework including traits coevolving across interacting lineages. Systematic Biology, 66: 551–568.

Mitov, V., Bartoszek, K., Asimomitis, G., and Stadler, T. 2020. Fast likelihood calculation for multivariate Gaussian phylogenetic models with shifts. Theoretical Population Biology, 131: 66– 78.

Mooers, A., Gascuel, O., Stadler, T., Li, H., and Steel, M. 2012. Branch lengths on birth–death trees and the expected loss of phylogenetic diversity. Systematic Biology, 61: 195–203.

Müller, N., Rasmussen, D., and Stadler, T. 2017. The structured coalescent and its approximations. Molecular Biology and Evolution, 34: 2970–2981.

Müller, K. 2020. here: A simpler way to find your files. R package version 1.0.1.

Neal, R. 2011. MCMC using Hamiltonian dynamics. In S. Brooks, A. Gelman, G. L. Jones, and X. L. Meng, editors, Handbook of Markov Chain Monte Carlo, pages 113–162. CRC Press, New York, NY.

Papieė, L. and Sandison, G. 1990. A diffusion model with loss of particles. Advances in Applied Probability, 22: 533–547.

Paradis, E. and Schliep, K. 2019. ape 5.0: an environment for modern phylogenetics and evolutionary analyses in R. Bioinformatics, 35: 526–528.

Preisler, H., Ager, A., and Wisdom, M. 2013. Analyzing animal movement patterns using potential functions. Ecosphere, 4: 1–13.

Pybus, O., Suchard, M., Lemey, P., Bernardin, F., Rambaut, A., Crawford, F., Gray, R., Arinaminpathy, N., Stramer, S., Busch, M., et al. 2012. Unifying the spatial epidemiology and molecular evolution of emerging epidemics. Proceedings of the National Academy of Sciences, U.S.A., 109: 15066–15071.

R Core Team 2024. R: A Language and Environment for Statistical Computing. R Foundation for Statistical Computing, Vienna, Austria.

Raghwani, J., Rambaut, A., Holmes, E., Hang, V. T., Hien, T. T., Farrar, J., Wills, B., Lennon, N., Birren, B., Henn, M., et al. 2011. Endemic dengue associated with the co-circulation of multiple viral lineages and localized density-dependent transmission. PLoS Pathogens, 7: e1002064.

Rakotomalala, M., Vrancken, B., Pinel-Galzi, A., Ramavovololona, P., Hebrard, E., Randrianangaly, J., Dellicour, S., Lemey, P., and Fargette, D. 2019. Comparing patterns and scales of plant virus phylogeography: Rice yellow mottle virus in Madagascar and in continental Africa. Virus Evolution, 5: vez023.

Russell, J., Hanks, E., Haran, M., and Hughes, D. 2018. A spatially varying stochastic differential equation model for animal movement. The Annals of Applied Statistics, 12: 1312–1331.

Sing, T., Sander, O., Beerenwinkel, N., and Lengauer, T. 2005. Rocr: visualizing classifier performance in r. Bioinformatics, 21(20): 7881.

Suchard, M., Lemey, P., Baele, G., Ayres, D. L., Drummond, A., and Rambaut, A. 2018. Bayesian phylogenetic and phylodynamic data integration using BEAST 1.10. Virus Evolution, 4: vey016.

Talbi, C., Lemey, P., Suchard, M., Abdelatif, E., Elharrak, M., Jalal, N., Faouzi, A., Echevarría, J.S.V.,, Rambaut, A., et al. 2010. Phylodynamics and human-mediated dispersal of a zoonotic virus. PLoS pathogens, 6: e1001166.

Taylor, J., Cumberland, W., and Sy, J. 1994. A stochastic model for analysis of longitudinal AIDS data. Journal of the American Statistical Association, 89: 727–736.

Tisseuil, C., Gryspeirt, A., Lancelot, R., Pioz, M., Liebhold, A., and Gilbert, M. 2016. Evaluating methods to quantify spatial variation in the velocity of biological invasions. Ecography, 39: 409–418.

Trovão, N., Baele, G., Vrancken, B., Bielejec, F., Suchard, M., Fargette, D., and Lemey, P. 2015. Host ecology determines the dispersal patterns of a plant virus. Virus evolution, 1: vev016.

van den Bosch, F., Hengeveld, R., and Metz, J. 1992. Analysing the velocity of animal range expansion. Journal of Biogeography, 19: 135–150.

Wang, L.-G., Lam, T. T.-Y., Xu, S., Dai, Z., Zhou, L., Feng, T., Guo, P., Dunn, C., Jones, B., Bradley, T., Zhu, H., Guan, Y., Jiang, Y., and Yu, G. 2019. Treeio: An R package for phylogenetic tree input and output with richly annotated and associated data. Molecular Biology and Evolution, 37: 599–603.

Wickham, H. 2016. ggplot2: Elegant Graphics for Data Analysis. Springer-Verlag New York.

Wilke, C. 2024. cowplot: Streamlined plot theme and plot annotations for ‘ggplot2’. R package version 1.1.3.

Wille, M., Geoghegan, J., and Holmes, E. 2021. How accurately can we assess zoonotic risk? PLoS Biology, 19: e3001135.

Wirtz, J. and Guindon, S. 2023. On the connections between the spatial Lambda-Fleming-Viot model and other processes for analysing geo-referenced genetic data. Theoretical Population Biology, 158: 139–149.

Wright, S. 1943. Isolation by distance. Genetics, 28: 114–138.

Zhu, H. 2024. kableExtra: Construct Complex Table with ‘kable’ and Pipe Syntax. R package version 1.4.0.

Zuckerkandl, E. and Pauling, L. 1965. Molecules as documents of evolutionary history. Journal of Theoretical Biology, 8: 357–366.

